# Single cell transcriptomics and developmental trajectories of murine cranial neural crest cell fate determination and cell cycle progression

**DOI:** 10.1101/2021.05.10.443503

**Authors:** Yu Ji, Shuwen Zhang, Kurt Reynolds, Ran Gu, Moira McMahon, Mohammad Islam, Yue Liu, Taylor Imai, Rebecca Donham, Huan Zhao, Ying Xu, Diana Burkart-Waco, Chengji J. Zhou

## Abstract

Cranial neural crest (NC) cells migrate long distances to populate the future craniofacial regions and give rise to various tissues, including facial cartilage, bones, connective tissues, and cranial nerves. However, the mechanism that drives the fate determination of cranial NC cells remains unclear. Using single-cell RNA sequencing combined genetic fate mapping, we reconstructed developmental trajectories of cranial NC cells, and traced their differentiation in mouse embryos. We identified four major cranial NC cell lineages at different status: pre-epithelial-mesenchymal transition, early migration, NC-derived mesenchymal cells, and neural lineage cells from embryonic days 9.5 to 12.5. During migration, the first cell fate determination separates cranial sensory ganglia, the second generates mesenchymal progenitors, and the third separates other neural lineage cells. We then focused on the early facial prominences that appear to be built by undifferentiated, fast-dividing NC cells that possess similar transcriptomic landscapes, which could be the drive for the facial developmental robustness. The post-migratory cranial NC cells exit the cell cycle around embryonic day 11.5 after facial shaping is completed and initiates further fate determination and differentiation processes. Our results demonstrate the transcriptomic landscapes during dynamic cell fate determination and cell cycle progression of cranial NC lineage cells and also suggest that the transcriptomic regulation of the balance between proliferation and differentiation of the post-migratory cranial NC cells can be a key for building up unique facial structures in vertebrates.

## Introduction

The craniofacial complex is one of the most diversified anatomical parts among vertebrates. Many studies have been focused on external factors like feeding behaviors that drive the selection of species-specific patterns of the craniofacial complex (Langenbach and van Eijden, 2001). Mutations that alter the molecular and cellular mechanisms of facial developmental processes could potentially provide the variation for selection, yet these mechanisms are still not well understood. The craniofacial complex comprise a cover of epithelial tissues, which are important signaling centers for craniofacial patterning (Hu and Marcucio, 2009), and mesenchymal tissues that build the substance of the complex and unique facial structures. Unlike the trunk mesenchyme that is derived from mesoderm, the facial mesenchyme mainly arises from cranial neural crest (NC) cells during development and evolution (Chai and Maxson, 2006, Dash and Trainor, 2020).

The NC cells are transiently existing multipotent stem cells and originated from the dorsal edge of the neural fold or neural tube during embryonic development (Ji et al., 2019, Crane and Trainor, 2006). After induction, the NC cells delaminate from the neuroectoderm and migrate long distances to generate numerous cell types (Shyamala et al., 2015). NC cells induced from the mid-diencephalon and hindbrain regions are named cranial or cephalic NC cells (Lumb et al., 2017). They migrate into the frontonasal prominence (FNP) and the first pharyngeal arch (Lumsden et al., 1991) to give rise to the paired lateral nasal prominences, the medial nasal prominences, the maxillary prominences, and the mandibular prominences, and eventually generate cartilage, bone, connective tissue, and other derivatives at the anterior region of the craniofacial complex and upper/lower jaws (Cordero et al., 2011, Chai and Maxson, 2006, La Noce et al., 2014).

When cranial NC cells arrive at the facial regions, the craniofacial complex is still very small. As the embryo develops, the NC cells undergo rapid proliferation to confer the correct shape and size of the craniofacial complex. Little is known about how the cranial NC cells generate different structures of the face. Some evidence shows that the fate of pre-migratory trunk NC cells can be predicted by the time they delaminate from the dorsal neural tube, as well as their ventral-dorsal position within the neural tube, suggesting there is an intrinsically programmed pre-patterning that regulates the early fate restrictions of NC cells (Nitzan et al., 2013, Krispin et al., 2010). However, unlike the trunk NC cells, which undergo EMT individually, the cranial NC cells delaminate from the neural tube and migrate as a coherent sheet of cells (Ji et al., 2019, Alfandari et al., 2010). Therefore, it is not known if defined NC-derived parts within the craniofacial complex arise from pre-determined regions of the dorsal neural tube. On the other hand, NC cells always interact with the mesoderm and ectoderm in the first pharyngeal arch to generate a fully functioning jaw (Baker and Bronner-Fraser, 2001). Signals, such as Shh, Wnt, Fgf and Bmp, from the epithelial signaling centers could guide the outgrowth of the frontonasal structures (Jheon and Schneider, 2009, Foppiano et al., 2007, Kasberg et al., 2013, Szabo-Rogers et al., 2008, Creuzet et al., 2004), indicating that the environment within the FNP may be critical for establishing the fate restrictions of NC cells. Therefore, whether the intrinsic or extrinsic cues that regulate cranial NC cells to commit to a restricted cell fate still needs further investigations.

In this study, we combined single-cell RNA sequencing (scRNA-seq) and lineage tracing to investigate the cell fate decisions involved in cranial NC lineage cells. We identified four major cranial NC cell lineages at different status and found that the cell fate determination that separates neural and mesenchymal cell lineages happens twice. We then focused on the NC- derived early mesenchymal cells right before the formation of the bones, cartilage, and other tissues in the face. We demonstrate that the differentiation of NC-derived mesenchymal cells initiates at a relatively late developmental stage, after the outgrowth of the craniofacial complex. Our results indicate that the initiation of the differentiation of NC-derived mesenchymal cells is coupled to cell cycle exit, indicating that instead of pre-patterned before migration, cell cycle regulation may play a more critical role in cranial NC cell fate determination and differentiation.

## Results

### A single-cell graph of NC developmental process in the murine embryos

We combined lineage tracing and scRNA-seq to address the spatiotemporal dynamics and investigate the cell fate decisions involved in NC cell differentiation after they migrate to their target regions (Fig. 1A). To genetically label NC cells and their progenies, we crossed Wnt1-Cre mice with the Rosa26-tdTomato/eGFP (Rosa26-mT/mG) reporter mouse line. The Wnt1-Cre-mediated recombination has been shown to occur in cranial and cardiac neural crest cells, midbrain, and developing neural tube (Tavares and Clouthier, 2015). Therefore, it can label these tissues and their progeny cells with green fluorescence. Since the cranial NC cells migrate to their target region around E8.5 and the generation of the facial prominences is completed by E12.5 (Ji et al., 2020), we collected eGFP positive cells from E9.5, E10.5, E11.5, and E12.5 embryos using fluorescence-activated cell sorting (Fig. 1B, C). Therefore, our data covered most of the circuital stages for forming, fusion, and merging craniofacial prominences (Ji et al., 2019). Then we performed 10X single-cell RNA sequencing using live eGFP positive cells. The sequencing detected roughly 3,382, 3,380, 3,012, and 2,465 genes per cell from E9.5, E10.5, E11.5, and E12.5 embryos, respectively (Fig. S1).

**Figure 1.**
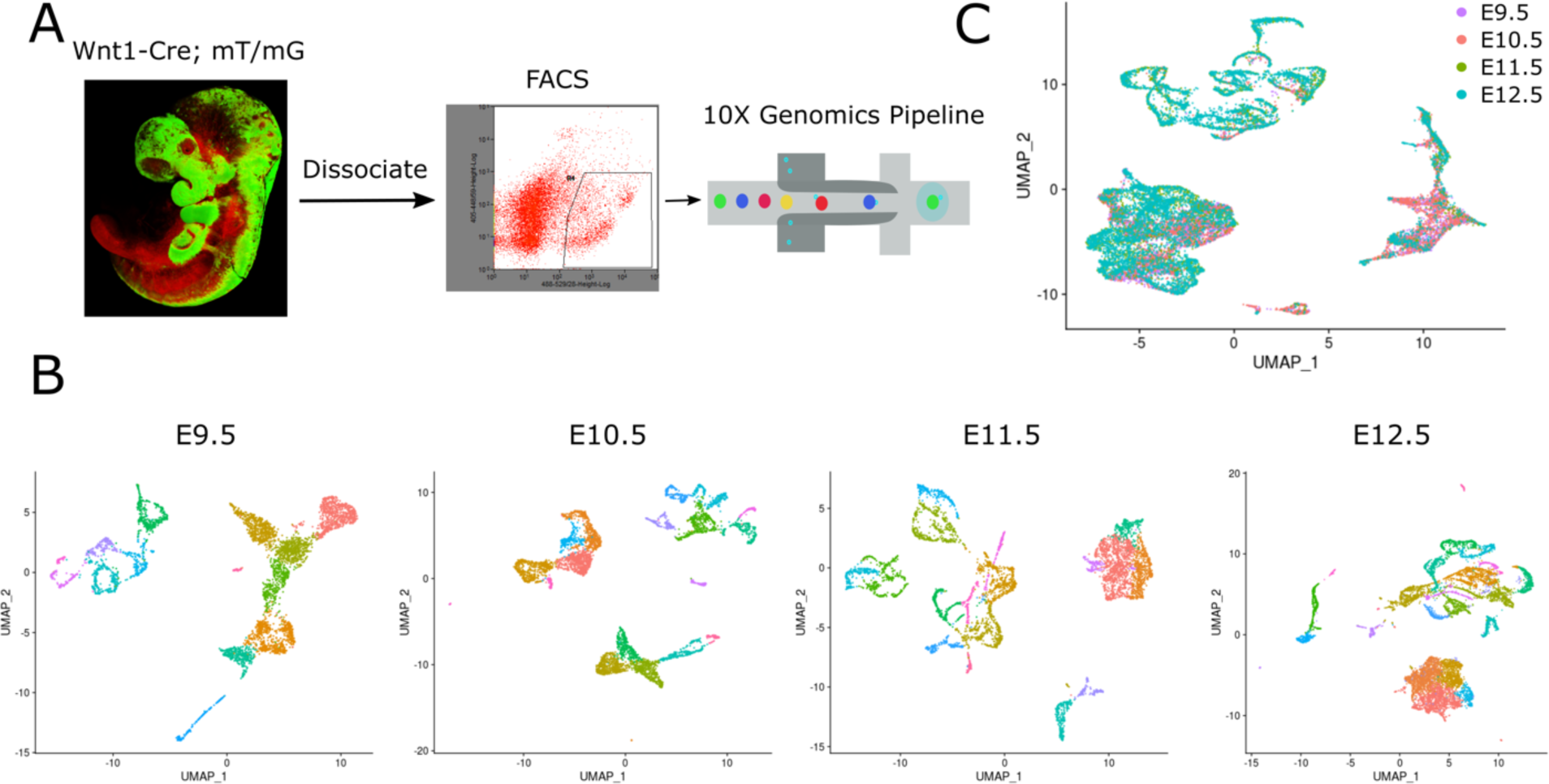
Single-cell transcriptional atlas of mouse NC cells. (A) Experimental workflow. NC cells were genetically labeled by Wnt1-Cre;Rosa26-mT/mG reporter. EGFP positive cells from E9.5, E10.5, E11.5, and E12.5 embryos were collected using fluorescent-activated cell sorting and performed 10X single-cell RNA sequencing. (B) UMAP plot for each time point. Non-NCCs were excluded from the following analysis based on expressed marker genes. (C) UMAP plot of 19,640 eGFP positive cells from four-time points. Cells are colored by time point.

In addition to the dorsal spinal cord, where the premigratory NC cells are located, the Wnt1-Cre is also expressed at the midlines of the midbrain, the caudal diencephalon, and the midbrain-hindbrain junction after the closure of the dorsal neural tube (Debbache et al., 2018). However, cells from the central nervous system can be distinguished from the NC-derived cells based on gene expression. We sorted out the non-NC cells based on their marker genes *in silico*. For example, the neural tube population expresses neural plate border specifier genes such as *Zic1*, *Zic3*, and *Pax3* (Soldatov et al., 2019), while cells from the developing midbrain, hindbrain, and cerebellum express *Wnt7b*, *Wnt8b*, *Hes5*, *Lhx1*, and *Lhx5* (Garda et al., 2002, Zhao et al., 2007). Moreover, epithelial cells (specifically expressing *Epcam*, *Krt8*, *Krt14*), endothelial cells (specifically expressing *Cdh5*), and red blood cells (specifically expressing *Hbb-y*, *Hba-x*, *Hba-a2*), which are small clusters containing only eGFP negative cells, were also removed from further analysis.

After removing the non-NC cells from our dataset, a total of 19,640 single-cell transcriptomes passed quality control for further analysis, including 3,620, 4,284, 4,302, and 7,434 cells from E9.5, E10.5, E11.5, and E12.5 embryos, respectively. Using Uniform Manifold Approximation and Projection (UMAP), we analyzed the transcriptional heterogeneity of NC-derived cells. When the data from all four time points were analyzed together, we found that the NC cells clustered into four major cell populations.

### Identification of the major NC-derived cell types

We identified the major NC-derived cell types based on the expression of known marker genes. The major NC-derived cell populations were identified as pre-EMT, which express *Wnt1* and *Wnt3a* (Ikeya et al., 1997), early migration, which express *Sox10* and *Foxd3* (Kirby and Hutson, 2010), NC-derived mesenchymal cells, which express *Twist1* and *Prrx2* (Soldatov et al., 2019), and the neural linage cells, which express *Tubb3* and *Elavl3* (Soldatov et al., 2019, Delile et al., 2019) (Fig. 2A, B, and Fig. S2, Table S1). Notably, a small cluster representing the cardiac neural crest cells that migrate to the outflow tract were identified based on their expression of *Tbx20* and *Acta2* (Singh et al., 2005). These results argue that our data captured a majority of known NC-derived cell types.

**Figure 2.**
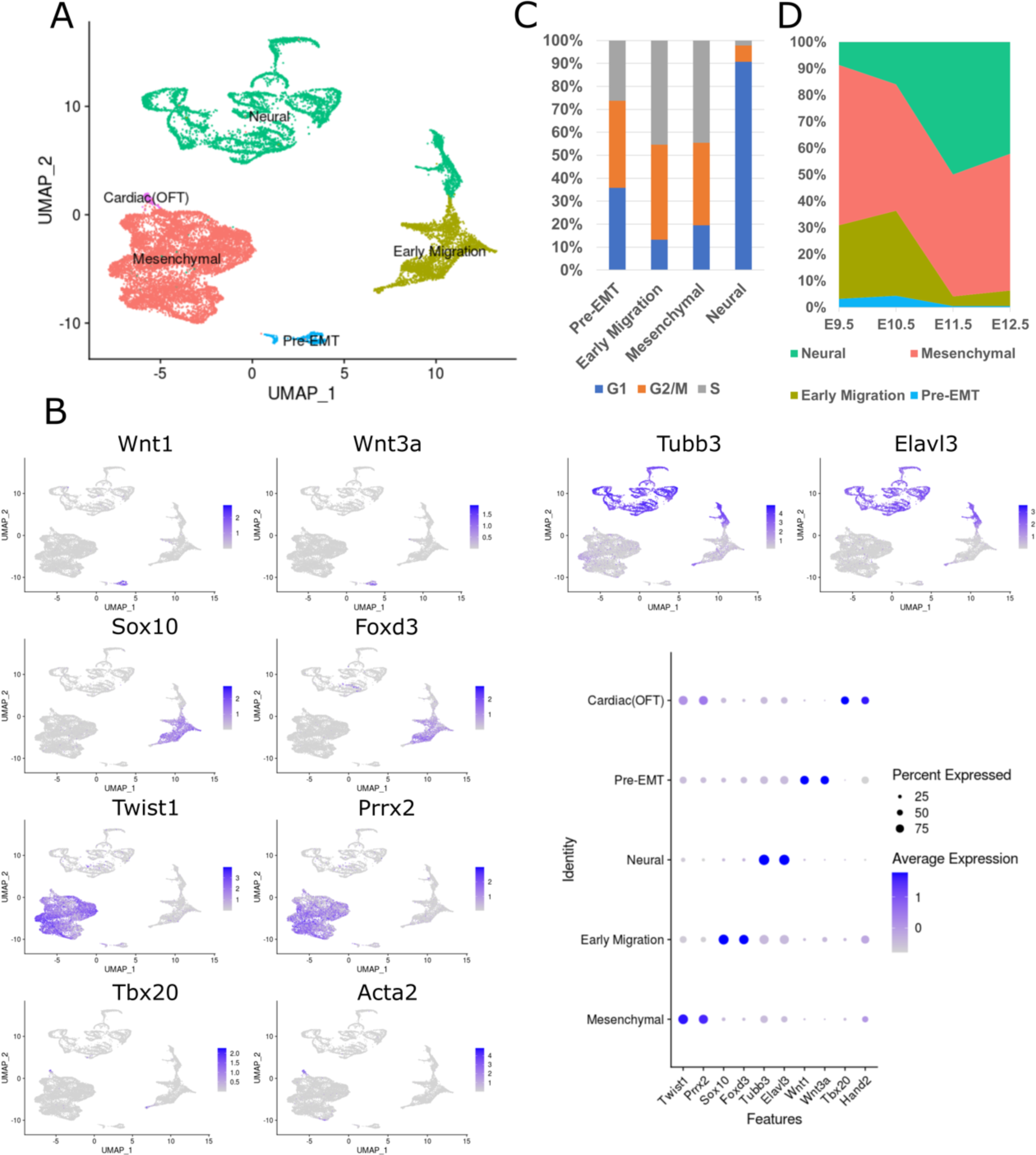
Single-cell RNA-seq identifies major NC-derived cell types. (A) UMAP plot of NC cells from four-time points. Cells are colored by major NC-derived cell types inferred from expressed marker genes. (B) Feature plots and dot plots of marker genes for each major cell population. (C) Histogram showing the fraction of different cell types in G1 (Blue), G2M (Orange), or S (Grey) phase. (D) The fraction of cell type at each time point, showing a decrease of pre-EMT and early migration NC cells after E11.5.

Previous studies showed that both pre-EMT and early migratory NC cells maintained multipotency in mice (Baggiolini et al., 2015). One important characteristic of multipotent stem cells is their ability to self-renew. Therefore, we analyzed the cell cycle states across the major cell populations using the Seurat scRNA-seq analysis pipeline (Fig. 2C, Fig. S3). Indeed, 64.2% of the pre-EMT and 86.7% of the early migratory neural crest cells are in the S or G2/M phase of the cell cycle, suggesting they are fast-dividing cells consistent with the hypothesis that they are multipotent progenitor cells. In contrast, only 9.3% of neural linage cells are at the S or G2/M phase, indicating that the neural linage is no longer actively dividing. Interestingly, our results showed that 80.5% of the mesenchymal cells are at the S or G2/M phase, indicating that most mesenchymal cells are also proliferating, the frequency of which was dynamic across the sampled time points (Fig. 2D). The frequency of pre-EMT and early migratory NC cells declined over time, while mesenchymal and neural lineage cells increased dramatically after E11.5. These results suggest that our data represent the developmental process that NC progenitor cells lose multipotency and differentiate into various fates.

### Reconstruction of developmental trajectories of NC cells

One of the benefits of scRNA-seq is that it allows mapping dynamic differentiation process by densely sampling cells at different stages, in our case E9.5, E10.5, E11.5, and E12.5. Therefore, these NC-derived cells sampled at different time points can be used to create a continually NC developmental trajectory. To reconstruct the progression that cells undergo during their differentiation from pre-EMT NC progenitor cells to fate determined neural or mesenchymal cells, we performed pseudotime analysis on all of the eGFP positive cells (Fig. 3A, B). The resulting lineage tree demonstrates the transcriptional changes associated with cell fate splits. After emigrating from the dorsal neural tube, the fate determination that separate neural and mesenchymal cell lineages happen twice (Fig. 3C). The first differentiation separates cranial sensory ganglia (expressing *Tlx2*, *Tlx3*, and *Neurog1*) (Logan et al., 1998) from progenitors of mesenchymal and the other neural linage, such as Schwann cell precursors (expressing *Pou3f1* and *Pu3f2*). The second fate determination generates the mesenchymal branch (Fig. 3D). Soldatov *et al*. also found that NC cell fate determination occurred through a progression of binary decisions, similar to our results (Soldatov et al., 2019).

**Figure 3.**
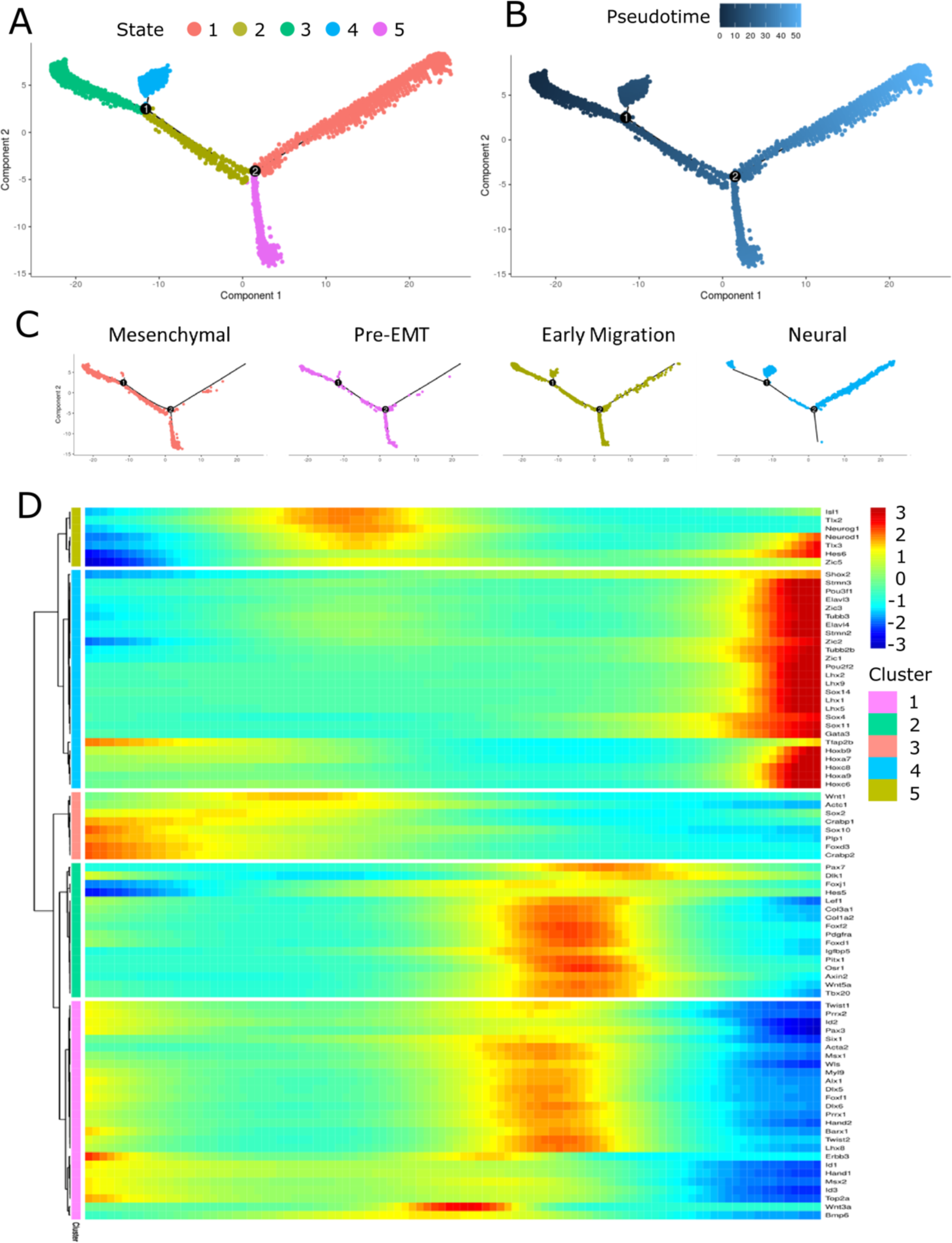
Developmental trajectories of NC cells. (A, B) Monocle pseudotime trajectory of NC cells from four-time points. Cells are colored by state (A) or pseudotime (B). (C) Monocle pseudotime trajectory showing the progression of pre-EMT, early migrating NC cells, mesenchymal, and neural lineages derived from NC cells. (D) Expression heatmap showing gene markers that link NC clusters to developmental states.

### Characterization of the mesenchymal cells at facial prominences

Once committed to a mesenchymal fate, cranial NC cells generate most of the bone and cartilage in the craniofacial regions. However, it remains unclear how post-migratory NC-derived mesenchymal cells are induced to differentiate into the specific craniofacial skeletal structures with the correct size and shape. To reveal the mechanisms that drive the fate determination of craniofacial mesenchymal cells, we re-clustered the 9,875 NC-derived mesenchymal cells collected from E9.5, E10.5, E11.5, and E12.5 embryos (Fig. 4A, B, Table S2). Using cluster-specific marker genes and wholemount *in situ* hybridization (WISH), we identified fifteen clusters associated with NC cells that migrate into the frontonasal prominences and the maxillary prominence of the first pharyngeal arch (Fig. 4C-E). For example, the aristaless-like homeobox *Alx* genes, *Alx1*, *Alx3,* and *Alx4*, are widely expressed in the midfacial complex at E10.5, and cells that highly express these genes are enriched in cluster m1 (mesenchymal 1), suggesting these clusters are from the midfacial prominences. By E11.5, *Alx1* and *Alx4* continue to be widely expressed in all three paired midfacial primordia, the lateral nasal prominence (LNP), the medial nasal prominence (MNP), and the maxillary prominence (MxP), suggesting that clusters m1, 3, 7, and 11 represent the mesenchymal cells at midfacial prominences. The expression of *Alx3* is limited to MNP, indicating that clusters m3 and 11 are the MNP. In contrast, *Barx1* is widely expressed in the mandibular prominence of the first pharyngeal arch, the second, third, fourth, and sixth pharyngeal arches at E10.5, suggesting clusters m2, 4, 5, 6, 8, 9, 10 are from the pharyngeal regions. At E11.5, a small part of ventral MxP also begins to express *Barx1*, and those cells are found in cluster m3. Together, using cluster-specific markers, we identified m1, 3, 7, and 11 as the mesenchymal cell populations of the midfacial prominences. Interestingly, cells from early developmental stages (E9.5 and E10.5) are enriched in cluster m1, while cells from late developmental stages (E11.5 and E12.5) are scattered in all four clusters, suggesting that the transcriptional heterogeneity for cells from E11.5 and E12.5 is more complicated compared to E9.5 and E10.5 (Fig. 4B).

**Figure 4.**
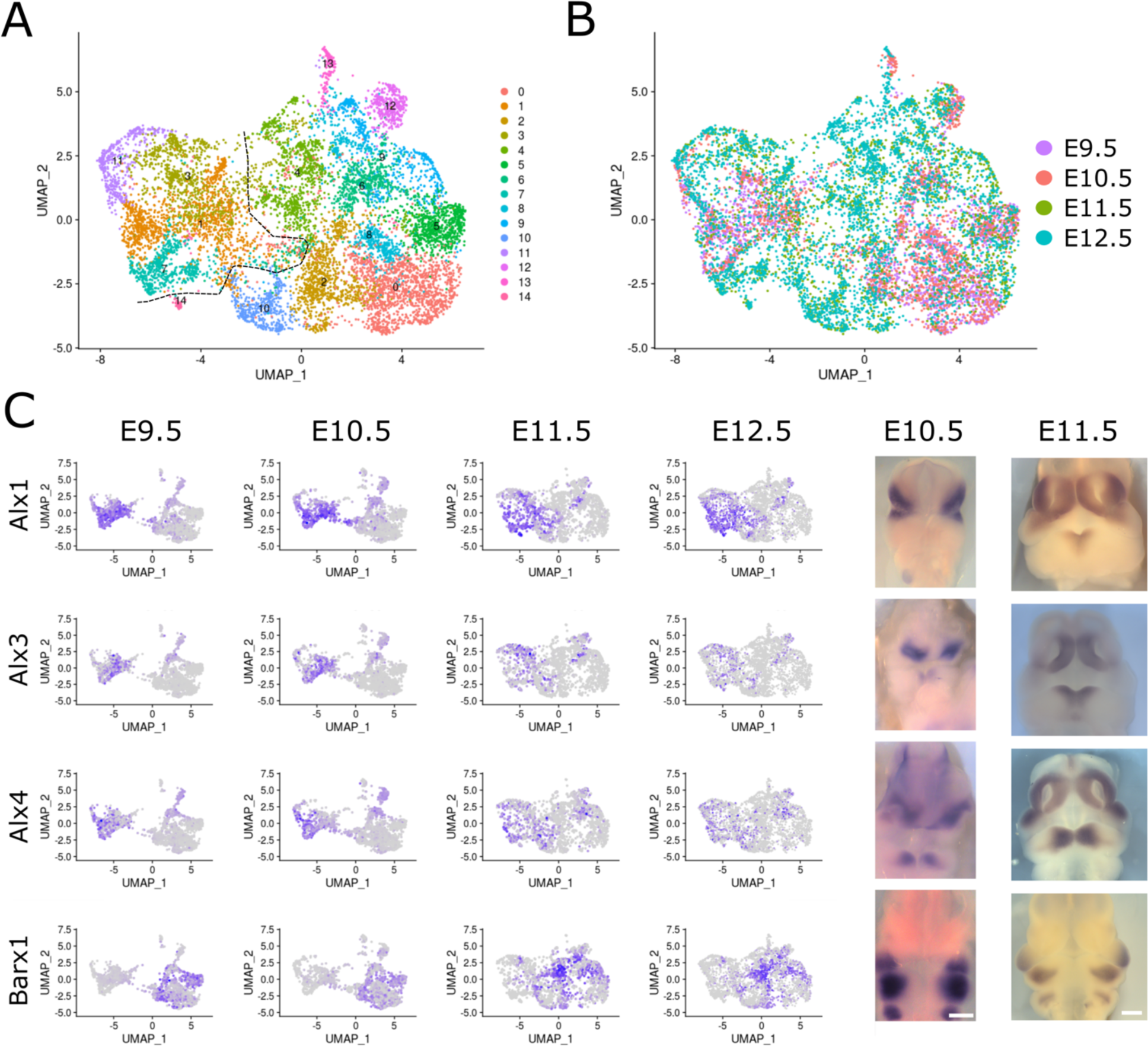
Single-cell RNA-seq identifies major NC-derived mesenchymal cells at facial primordia. (A, B) UMAP plot of 9,875 NC-derived mesenchymal cells showing 15 different cell populations. Cells are colored by mesenchymal clusters (A) or time point (B). (C) Identifying the mesenchymal clusters using wholemount *in situ* hybridization. Left panels, feature plots for marker genes. Right panels, wholemount *in situ* hybridization results for indicated genes at E10.5 and E11.5. Scale bar: 0.5mm

### The differentiation of NC-derived craniofacial mesenchymal cells starts at a relatively late developmental stage

We next performed a pseudotime analysis of cells from clusters m1, 3, 7, and 11 using Monocle 2 to reconstruct the development processes (Trapnell et al., 2014). Monocle 2 performed an unsupervised analysis to order the cells and reconstructed a tree-like trajectory, beginning with E9.5 and E10.5 cells and ending with E11.5 and E12.5 cells (Fig. 5A, B). Notably, cells in the “root” branch are mostly from E9.5 and E10.5 embryos. By contrast, cells from E11.5 and E12.5 embryos are spread out in the other four states, indicating that the differentiation of craniofacial mesenchymal cells is likely to initiate at a later developmental stage (Fig. 5C).

**Figure 5.**
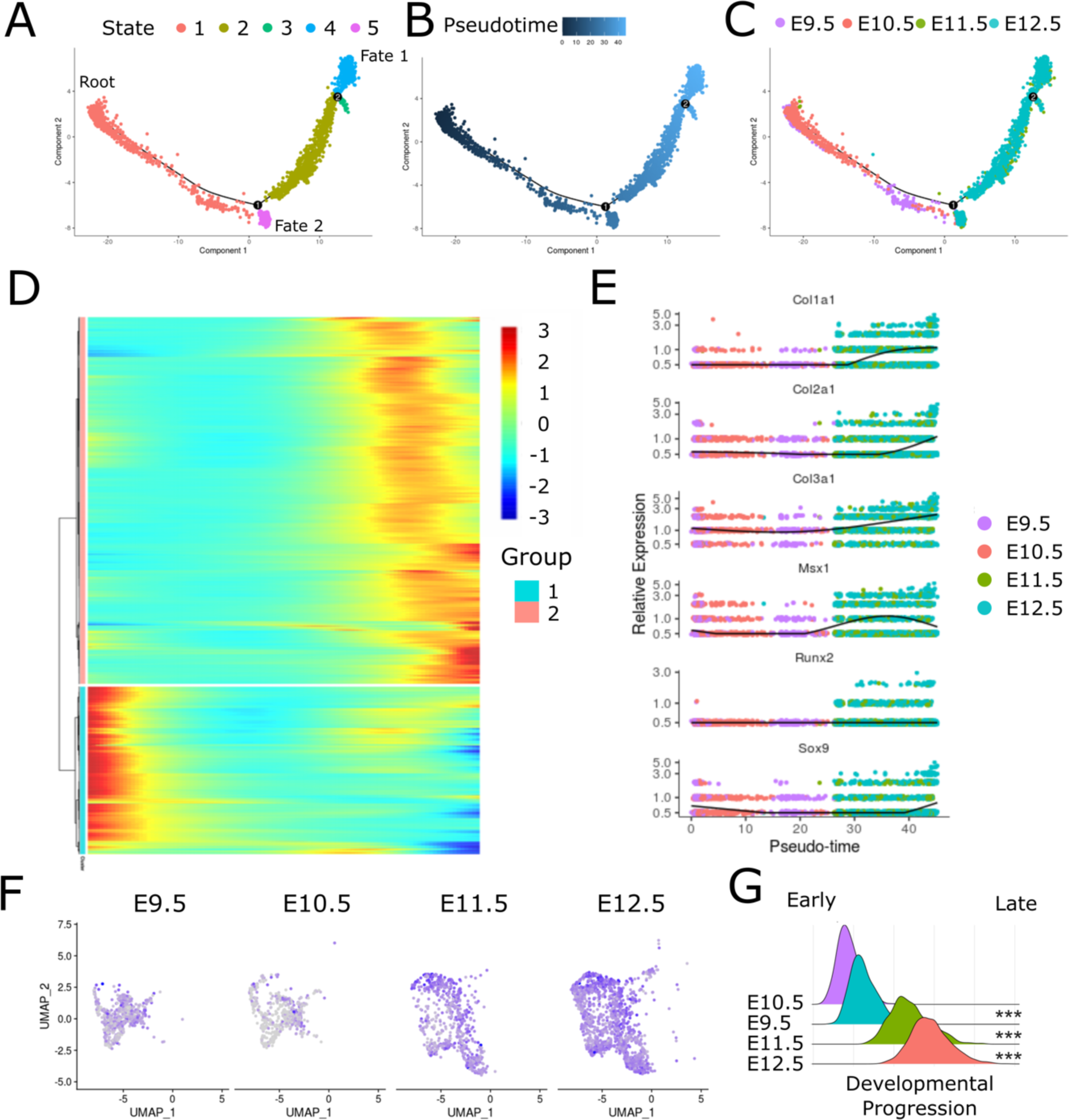
The differentiation of NC-derived craniofacial mesenchymal cells starts at late developmental stages. (A-C) Monocle pseudotime trajectory of NC-derived craniofacial mesenchymal cells from four-time points. Cells are colored by state (A), pseudotime (B), or time point (C). (D) Global analysis of gene expression along the trajectory identified over 10,000 genes exhibits temporal expression patterns. (E) Pseudotime kinetics show the expression of known gene markers of the cell cycle and differentiation process. (F, G) Feature plot (F) and histograms (G) showing the developmental progression increased after E11.5 using a developmental progression score for each cell. Wilcoxon rank sum test was used to compare samples from each time point with E9.5 sample. *** = p< e^-14^

The transcriptional complexity of NC lineage cells decreases with developmental progression (Jean-Baptiste et al., 2019, Saunders et al., 2019, Gulati et al., 2020). Dorrity et al. developed a mathematical model to calculate the developmental progression of a single cell based on the number of genes that were detected per cell (Dorrity, 2020). To test whether the differentiation of craniofacial mesenchymal cells starts after E11.5, we measured the developmental progression of craniofacial mesenchymal cells (Fig. 5F, G). The result shows that the transcriptional complexity decreases in E11.5 and E12.5 embryos, leading to a higher developmental progression score, while E9.5 and E10.5 embryos remain at early developmental stages (Fig. 5F, G). In addition, we identified that more than 10,000 genes that exhibit temporal expression patterns, which fall into two distinct gene groups (Fig. 5D, Table S3). One group of genes are highly expressed in the early development stages (E9.5 and E10.5). The other group of genes is highly expressed at the late development stages (E11.5 and E12.5) and includes genes like *Msx1* and *Col1a1* that are critical for craniofacial development (Fig. 5E), suggesting the ossification of craniofacial mesenchymal cells initiates at E11.5.

To test whether the ossification of NC-derived craniofacial mesenchymal cells starts at E11.5, we examined the expression of genes regulating bone formation in cells at different time points along the developmental trajectory. The ossification of undifferentiated mesenchymal cells into bone cells begins with the formation of osteoprogenitors. This step is regulated by master transcription factors such as *Sox9*, *Runx2,* and *Msx1* (Javed et al., 2010). The expression of these genes increased after E11.5 along the trajectory (Fig. 5E). During the next stage of osteoblast development, the cells start to express genes like collagen and fibronectin genes (Rutkovskiy et al., 2016), which are known to be critical for the arrest of cell motility during the osteoblast-to-osteocyte transition (Shiflett et al., 2019). In our data, the expression of *Col1a1*, *Col2a1*, and *Col3a1* is increased after the first branch point along the trajectory (Fig. 5E), suggesting the ossification of craniofacial mesenchymal cells initiates after E11.5. Therefore, although NC-derived mesenchymal cells populate at the craniofacial complex and give rise to different prominences as early as E9.5, they remain at an undifferentiated, homogeny stage. The differentiation of NC-derived mesenchymal cells into different bones in the facial region does not initiate until E11.5, suggesting the fate of cranial NC cells might not be intrinsically programmed but is acquired from the environment.

### The initiation of the 2ossification is coupled to cell cycle progression

To reveal the mechanism that triggers the differentiation of cranial NC-derived mesenchymal cells, we performed branch-dependent expression analysis at the first branch point (Fig. 6A, Table S4). The results showed that one group (Group 1 in Fig. 6A) of genes is highly expressed in the root branch, and the expression of these genes significantly decreased in both branches, especially in cells at fate 1. Gene ontology (GO) enrichment analysis suggests that these genes are involved in cell cycle regulation, chromosome segregation, and microtubule cytoskeleton organization. In addition, many genes that inhibit the differentiation process are also downregulated in both branches (Fig. 6B). For example, the inhibitor of DNA binding and cell differentiation protein (Id1) has been shown to be expressed in embryonic and somatic stem cells and sustains the stemness of these cells through inhibition of differentiation (Jankovic et al., 2007, Nakashima et al., 2001, Nam and Benezra, 2009, Ying et al., 2003). *Id1* expression was downregulated in both cell fates. *Stub1*, encoding an E3 ligase that negatively regulates ossification by inducing the degradation of Runx2, is also down-regulated in both branches (Li et al., 2008). These results suggest that at E11.5, the NC-derived craniofacial mesenchymal cells adopt a ready-to-differentiate state.

**Figure 6.**
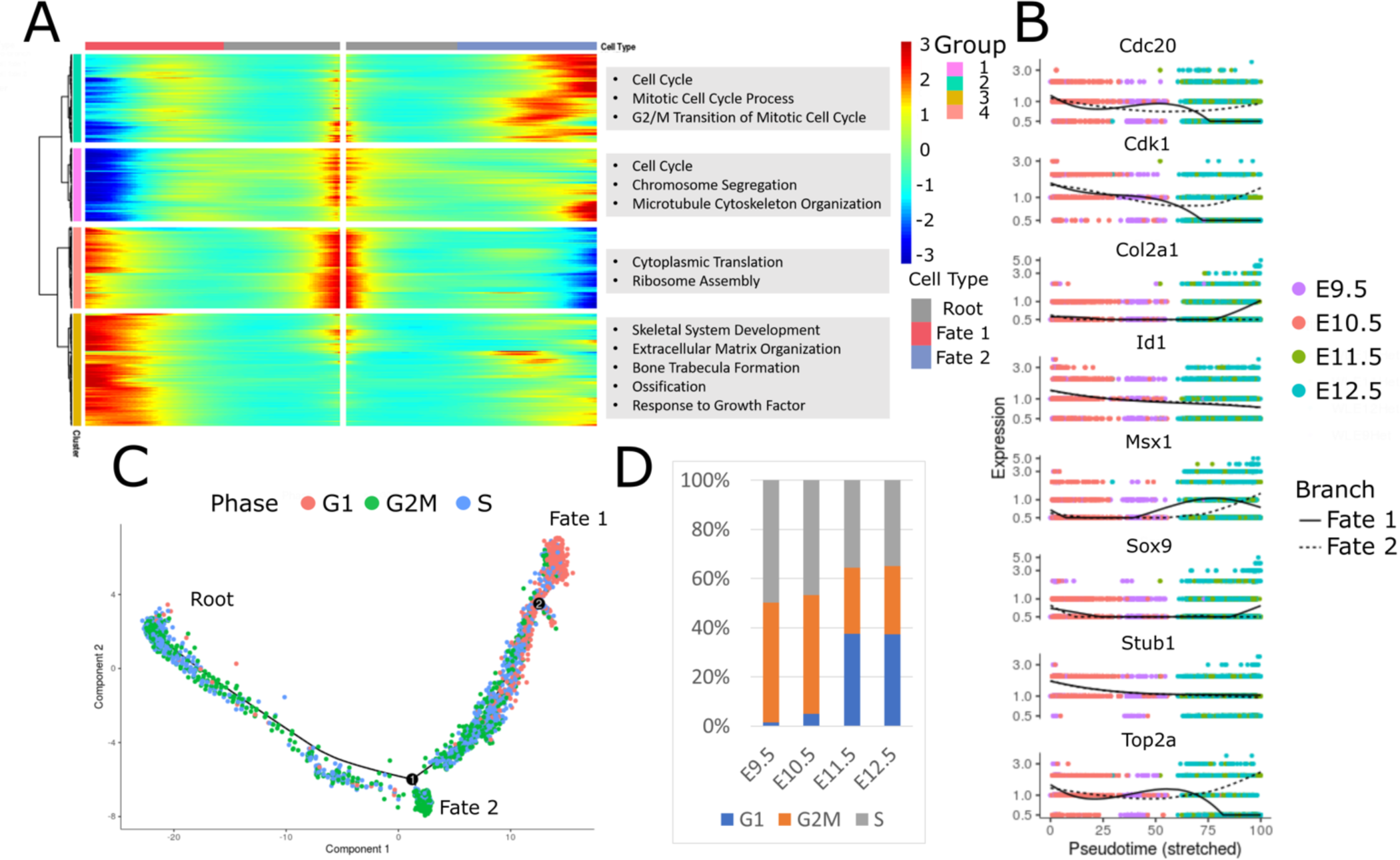
Branch 1 is a decision-making point governing whether a cell exits the cell cycle (A) Expression heatmap showing genes expressed in a branch 1 -dependent manner. Pre-branch refers to the cells before the branch. Gene ontology (GO) enrichment analysis for genes from each cell fate is shown (Right panel). (B) Pseudotime kinetics show the expression of known gene markers of the cell cycle and differentiation process from the root of the trajectory to fate 1 (solid line) or fate 2 (dotted line). (C) Pseudotime trajectory of craniofacial mesenchymal cells from four-time points. Cells are colored by the cell cycle phase. (D) Histogram showing the fraction of cells in G1 (Blue), G2M (Orange), or S (Grey) phase at each time point.

Another group of genes (Group 3 in Fig. 6A) has been shown to be highly expressed in cells at one branch (fate 1 in Fig. 5A and Fig. 6A). Functional analysis of genes in this group suggests that they are involved in skeletal development, extracellular matrix organization, and ossification (Fig. 6A). For example, bone-specific genes, such as collagens, *Sox9*, and *Msx1*, are specifically up-regulated in fate 1. On the other hand, cell cycle genes, including *Top2a*, *Cdk1*, *Cdc20*, and *Rrs1*, are upregulated in fate 2 (Fig. 6B). Moreover, 98.5% and 95% of cells from E9.5 and E10.5 embryos are dividing cells, while only 62.5% and 62.7% of cells from E11.5 and E12.5 embryos are dividing cells (Fig. 6C, D), indicating that a group of cells has exited the cell cycle by E11.5. Given that the first branch point of the trajectory separates cells before or after E11.5, our results indicate that this branch point might be a decision-making point for a population of cells to exit the cell cycle and start to differentiate.

To address the reproducibility of our analysis, we isolated post-migratory NC cells labeled by Sox10-Cre;Rosa26-mT/mG reporter from E11.5 embryos, and performed scRNA-seq analysis (Hari et al., 2012). Indeed, the facial mesenchymal cells from the Sox10-Cre dataset generated the same four clusters with a similar percentage of cells in each cluster as the Wnt1- Cre dataset. Moreover, the expression level of all genes, the patterns of marker genes were also similar between Wnt1-Cre and Sox10-Cre datasets (Fig. 7). Therefore, our analysis revealed highly reproducible NC-derived cell populations associated with facial prominences, which allows us to combine Wnt1-Cre and Sox10-Cre datasets for further analysis.

**Figure 7.**
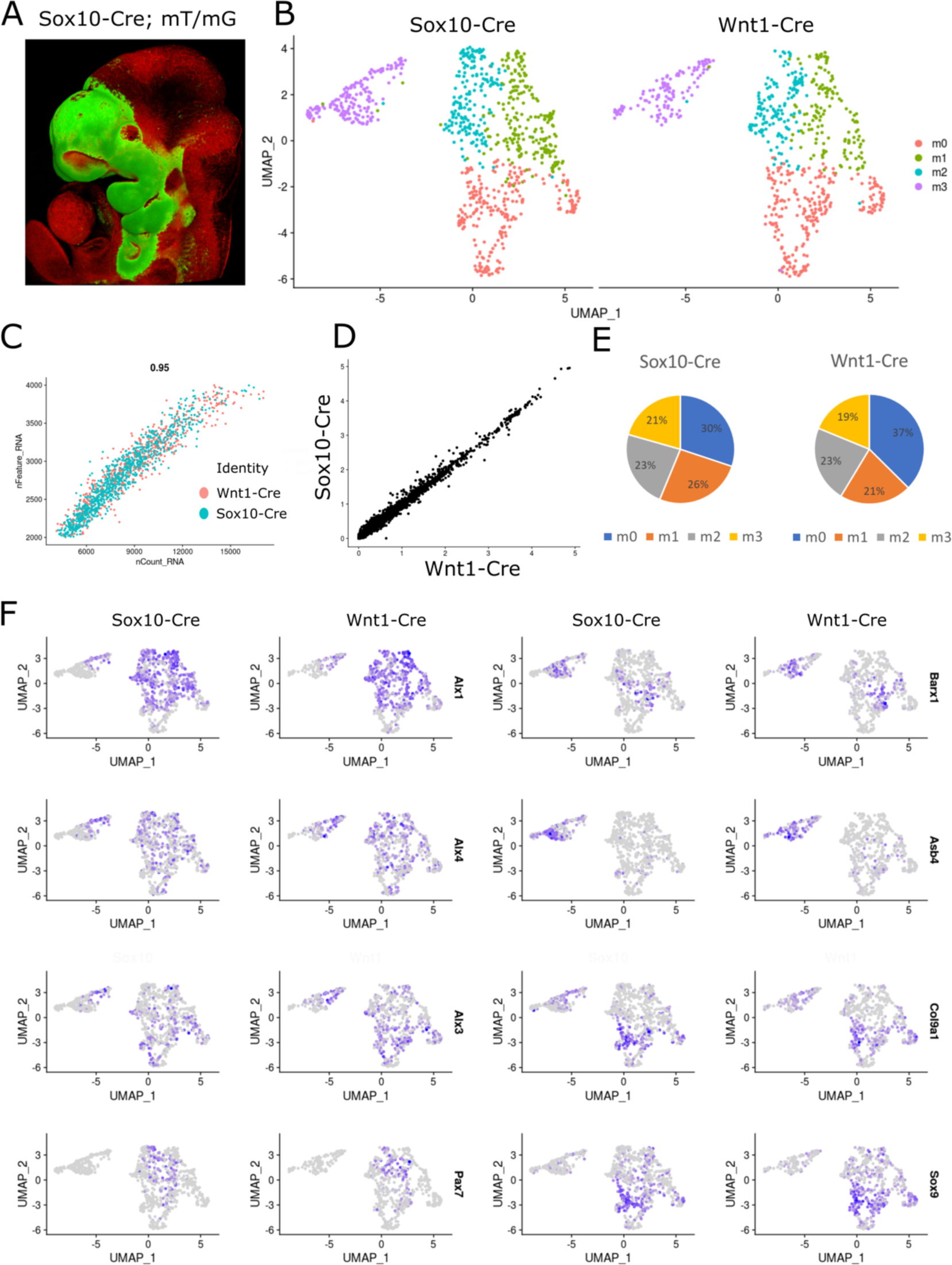
High reproducibility between Wnt1-Cre and Sox10-Cre scRNA-seq dataset at E11.5. (A) Confocal imaging shows Sox10-Cre marks migratory NC cells and their progeny. (B) UMAP plot shows the transcriptional dynamics are similar between Wnt1-Cre and Sox10-Cre scRNA-seq dataset. (C) Distributions of the number of UMIs and genes detected in the two datasets is similar. (D) Scatter plot shows the average expression of each gene in each cell is similar between the two datasets. (E) Pie charts show that the percentages of each cell type are similar. (F) Feature plots of marker genes show the expression patterns of these genes are similar between the two datasets.

The facial mesenchymal cells from E11.5 embryos were populated into four clusters. Based on marker genes and published literature (Li et al., 2019), we found that m0, m1, and m2 contain cells from MNP and LNP (expressing *Alx1*, *Alx3*, *Alx4*, and *Pax7*), and m3 contains cells from MxP (expressing *Barx1* and *Asb4*) (Figs. 7F, 8A, B, Table S5). Cell cycle analysis indicated that 83.2% of cells in m2 and 67.7% in m3 are actively dividing cells while only 46.1% m0 and 44.9% m1 cells are undergoing mitosis (Fig. 8C, D). Further analysis revealed that cells in m0 and m1 exhibit more transcriptional complexity than cells in m2 and m3, indicating that m2 and m3 might be at an earlier developmental stage than m0 and m1 (Fig. 8E). During mouse facial development, the morphogenesis of the LNP- and MNP-derived structures progresses from E10.5 to E12.5. However, MxP cells continue to grow until E15.5 to give rise to the secondary palate (Ji et al., 2020). This is consistent with our results that, at E11.5, LNP and MNP cells (m0, m1, and m2) were grouped into three clusters at different differential stages while most of the MxP cells (m3) are still at an early developmental stage.

**Figure 8.**
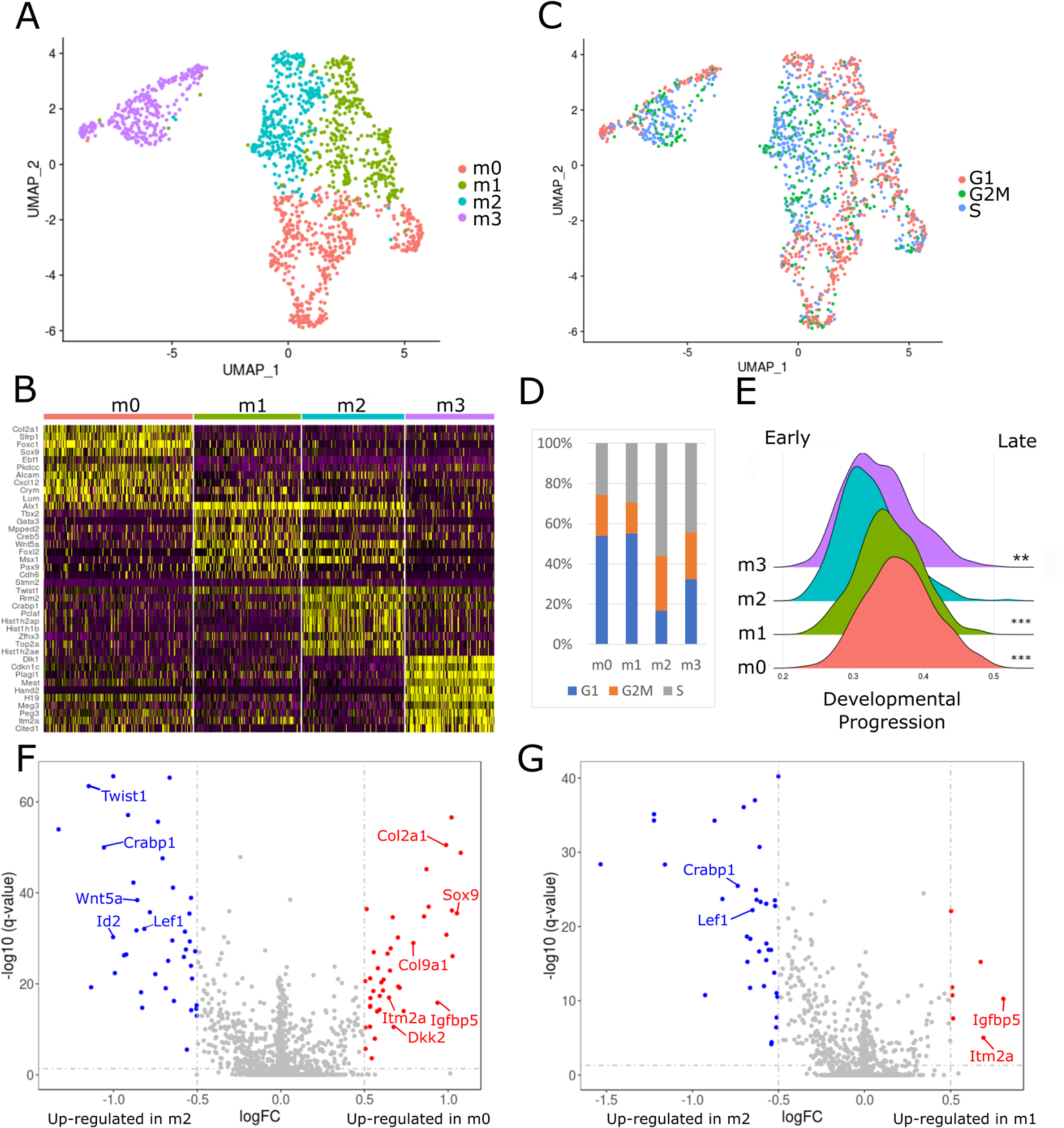
The development of craniofacial mesenchymal cells is associated with cell cycle progression at E11.5. (A) UMAP plot of NC-derived craniofacial mesenchymal cells from E11.5 embryos. (B) Gene expression heatmap of top 10 marker genes for each cluster. (C) Segregation of craniofacial mesenchymal cells by cell cycle phase. (D) Quantification of the proportion of different cell types in G1 (Blue), G2M (Orange), or S (Grey) phase. (E) Histograms showing the developmental progression score for each cell type. Wilcoxon rank sum test was used to compare cells from each cluster with m2. *** = p< e^-14^, ** = p< e^-3^ (F, G) Volcano plots showing differentially expressed genes between m2 and m0 (F) or m2 and m1 (G). Highly expressed genes (avg_logFC > 0.5, P adj < 0.05) are colored in blue (in m2) or red (in m0 or m1) dots, respectively.

To further reveal the possible mechanisms that drive the differentiation of LNP and MNP cells, we analyzed the differentially expressed gene expression between m2 and m0 (Table S6) and m2 and m1 (Table S7). As a result, both *Twist1* and *Id2*, known to be required for cell proliferation during the early osteoblast differentiation stage (Javed et al., 2010, Sakata-Goto et al., 2012), were expressed at a higher level in m2. In contrast, many genes involved in cartilage formation, such as *Sox9*, *Col2a1*, *Itm2a*, *Igfbp5*, and *Col9a1*, are expressed at a higher level in m0 than in m2, suggesting cells in m0 are chondroprogenitors (Fig. 8F). Although cells in m1 do not have many highly expressed genes compared to m2, *Igfbp5* and *Itm2a* were found to also highly express in m1. *Igfbp5* has been shown to stimulate bone cell growth (Miyakoshi et al., 2001). Studies in mice also showed that *Itm2a was* involved in osteogenic differentiation (Tuckermann et al., 2000). These results indicate that m1 might represent a transition stage between mesenchymal stem cells and chondroprogenitors. Moreover, we also found that *Wnt5a*, *Lef1*, and *Crabp1* were highly expressed in m2. In contrast, a WNT antagonist gene *Dkk2* was highly expressed in m0, suggesting Wnt and retinoic acid signaling might be essential for maintaining the self-renewal of mesenchymal stem cells.

## Discussion

The facial region is mainly comprised of NC-derived cells. However, how cranial NC cells develop into the facial structures is still not entirely clear. We addressed this question by tracing the lineage of NC cells in mouse embryos. Using the single-cell transcriptomic data, we described a spatiotemporal molecular specification tree of post-migratory cranial NC cells, showing the fate determination process of NC-derived mesenchymal cells at the facial prominences. Our study indicates that the differentiation of NC-derived craniofacial mesenchymal cells initiates as late as E11.5, and the differentiation is coupled with the exit of the cell cycle.

In mice, the cranial NC cells migrate to the anterior of the embryos as early as E8.5 and build the frontonasal prominence (FNP) at E9.25. By E10.25, the FNP gives rise to the paired MNPs and LNPs. The MNPs continue to fuse with the MxP at E11.25, generating the upper jaw (Everson et al., 2018). Eventually, the cranial NC cells generate most of the bone and cartilage in the craniofacial region. However, how the fate of each NC cell is determined is unclear. One hypothesis was that NC cells contain an intrinsically programmed molecular facial patterning “blueprint” when they delaminate from the neural tube. However, our single-cell transcriptome data shows that at E9.5 and E10.5, NC-derived mesenchymal cells in different facial prominences are similar to each other at the transcriptomic level, regardless of where they come from. At a later developmental stage, E11.5, the cells exhibit more transcriptional heterogeneity, suggesting that the fates of different cell populations are beginning to diverge. Reconstruction of a lineage trajectory of cranial NC lineage cells also indicates the differentiation of NC cells initiates at E11.5. Therefore, we propose that cranial NC cells maintain their differentiation potential until the morphology of the face is shaped. In support of our new hypothesis, Kaucka *et al*. showed that the shape of the face is mainly formed by local cellular divisions (Kaucka et al., 2016). The authors also found that the proliferation activity of NC cells resulted in cellular mixing in the facial tissue. They proposed that the fact that progenies of several NC cells locally mixed might guarantee the developmental robustness of facial complex. Our results support this hypothesis because we found that fast-dividing NC cells maintain their stemness. Therefore, hypothetically, mutations that happen in an NC cell can be counterbalanced by unaffected neighbor undifferentiated NC cells.

Terminal differentiation of many multipotent cells such as neural stem cells is associated with cell cycle exiting (Soufi and Dalton, 2016, Hardwick and Philpott, 2014). Our results showed that the NC-derived mesenchymal stem cell differentiation is also cell cycle-dependent. The molecular mechanisms that underline this phenomenon still need further study. One possible mechanism is the epigenetic landscape at the developmental genes that might change in the G1 phase. In pluripotent stem cells, most developmental genes are H3K4 and H3K27 trimethylated near their transcription start sites to be silenced by a polycomb-dependent mechanism (Bernstein et al., 2006). However, transcriptional leakiness of developmental genes could happen at the G1 phase, making the G1 phase an opportunity for differentiation (Singh et al., 2013). Consistent with this hypothesis, ablation of the plycomb complex-associated methyltransferase gene *Ezh2* in NC cells causes various facial deformities (Kim et al., 2018, Schwarz et al., 2014). Also, BMP signaling has been shown to be required for guiding the outgrowth of facial prominences (Graf et al., 2016). G1-specific cyclin-dependent protein kinases (CDKs) have been shown to target transcription factors like Smad2/3, leading to the expression of their target genes in the G1 phase (Kim et al., 2018, Pauklin and Vallier, 2013). Cell cycle regulation has been known to play an important role in many development processes of NC cells. For example, trunk NC cells delaminate from the dorsal neural tube only in the S phase (Burstyn-Cohen and Kalcheim, 2002). *In vivo* studies also revealed the dividing activity of cranial NC cells is increased as they migrated into the branchial arches (Ridenour et al., 2014). These cells continue dividing to form the facial complex with the correct shape and size (Kaucka et al., 2016). Our study shows that at a relatively late developmental stage (E11.5), after building the morphology of the craniofacial structures, the NC cells exit the cell cycle and start to differentiate. This may indicate that the variety of craniofacial shapes and functions in different species might be regulated by the rates of proliferation and the time exiting the cell cycle.

The fact that cranial NC-derived mesenchymal cells keep rapidly dividing until E11.5 suggests some signals promoting fast NC cell proliferation. Comparing the gene expression of fast and slow dividing cells from MNPs and LNPs reveals that *Wnt5a* is highly expressed in the fast-dividing cells, which has been shown to orient the direction of cell division and outgrowth of facial structures (Kaucka et al., 2016). Our results show that NC-derived mesenchymal proliferation in the facial primordia is, at least partly, regulated by *Wnt5a*. This conclusion is supported by the fact that knocking out and overexpressing *Wnt5a* causes facial outgrowth deficiency (van Amerongen et al., 2012, Bakker et al., 2012, Ho et al., 2012), suggesting the gradient of *Wnt5a* needs to be precisely regulated within the facial primordia for developing a face with correct shape and size.

Our data highlight the similarity between cranial NC cells at early developmental stages at a single cell transcriptomic level, which could be the reason for the facial developmental robustness. Additionally, our data also reveal that NC cell differentiation is associated with the exiting of the cell cycle, and the regulation of cell proliferation might be a key step in the evolution of various craniofacial shapes and functions in vertebrates.

## Materials and methods

### Mouse strains and genetic fate mapping

All animal work was approved and permitted by the UC Davis Animal Care and Use Committee and conducted according to the NIH guidelines. NC cell-specific genetic tracing mouse lines, Wnt1-Cre and Sox10-Cre lines were previously described (Lewis et al., 2013, Chen et al., 2017, Hari et al., 2012). Both Wnt1-Cre and Sox10-Cre strains were crossed with the Rosa26- tdTomato/eGFP (Rosa26-mT/mG) reporter line to label NC cells with eGFP (Muzumdar et al., 2007). All the mouse strains were ordered from the Jackson Laboratory (stock numbers 022137, 025807, and 007676). Pregnant, timed-mated mice were euthanized with overdosed isoflurane (SAS, PIR001325-EA) prior to cesarean section. The day of conception was designated embryonic day 0.5 (E0.5). For genetic fate mapping, Wnt1-Cre;Rosa26-mT/mG and Sox10- Cre;Rosa26-mT/mG embryos were sampled at E10.5. The wholemount imaging was performed with Nikon A1 confocal laser microscope. Basic image processing and analysis were performed using NIS-Elements C software.

### Wholemount *in situ* hybridization

E10.5 and E11.5 mouse embryo cDNA libraries were used to clone fragments of the coding sequence of mouse *Alx1*, *Alx3*, *Alx4*, and *Barx1*. The following primers were used: *Alx1*: forward primer 5′-GCGAGAAGTTTGCCCTGA-3′, reverse primer 5′-AAATGCGTGTCCGTTGGT-3′, *Alx3*: forward primer 5′-CTGTCTCATGTCTCCAGAGGG-3′, reverse primer 5′- TGTAGACTAGCACAGGGCAGAA-3′, *Alx4*: forward primer 5′-CCATCCTGGATTGGCAAC-3′, reverse primer 5′-GGGGGCCTGACTTTGACT-3′, *Barx1*: forward primer 5′- AGACAATTAAGGGCCAGACAAG-3′, reverse primer 5′-GTCCCCCACTGTGTCATAAAAT-3′. The embryos were collected at E10.5 and E11.5 and fixed in 4% PFA overnight at 4 °C. As previously described, wholemount *in situ* hybridization was performed with digoxigenin-labeled antisense RNA probes (Volker et al., 2012). Briefly, embryos were digested with proteinase K (1:1000) for 6 min (for E10.5 embryos) or 20 min (for E11.5 embryos) and refixed in 4% PFA/0.25% glutaraldehyde for 20 min at room temperature. After 60 min of prehybridization at 67 °C, the embryos were hybridized with probes overnight at 67 °C. Embryos were incubated with 1:4,000 diluted alkaline phosphatase-conjugated anti-digoxigenin antibody (Roche, 11093274910) overnight at 4 °C. Alkaline phosphatase activity was detected using NBT/BCIP (Sigma, N6876 and 10760994001).

### Single-cell RNA-sequencing analyses

Mouse embryos with Rosa26-mT/mG reporter were collected at E9.5, E10.5, E11.5, and E12.5 in ice-cold PBS and then dissociated using a cold-active protease (CAP) protocol (Adam et al., 2017). Briefly, embryos were incubated to 125 µl of cold protease solution [1.25 mg/ml *Bacillus Licheniformis* protease (Creative Enzymes, NATE0633) and 125 U/ml DNAseI (ThermoFisher, EN0521) in Dulbecco’s phosphate-buffered saline (DPBS) with calcium and magnesium] in 4 °C with trituration using a 1 ml pipet (10 s every 3 min). After 9, 12, 21, and 30 min incubation for E9.5, E10.5, E11.5, and E12.5 embryos, respectively, 1 ml ice-cold PBS with 15% fetal bovine serum (PBS/FBS) was added to the single-cell suspension. Cells were passed through a 35 µM cell strainer (Falcon, 352235). Cells were pelleted by 1200 g centrifuge for 5 min at 4 °C and re- suspended in 1 ml PBS with 1% FBS. The PBS/FBS wash was repeated one more time. 1000nM DAPI was added to the cell suspension to label dead cells. Fluorescent-activated cell sorting was performed to collect eGFP positive and DAPI negative cells. The cell concentration was adjusted to approximately 500 cells/ µl for 10x Genomics’ single-cell RNA-seq.

The Cell Ranger Single Cell software (http://10xgenomics.com/) was used to align reads and generate feature-barcode matrices. Cell clusters and marker genes were identified using Seurat_ 3.2.0 (Butler et al., 2018, Stuart et al., 2019). Initial cell filtering selected cells that expressed >2000 reads and contained <10% mitochondrial genes. Normalization was performed by the “NormalizeData” function in Seurat. The “FindVariableFeatures” function was used to calculate a subset of highly variable features (10000 genes) for future analysis. We used the “CellCycleScoring” function in Seurat to score the cell cycle phase of every cell (Nestorowa et al., 2016). The cell that highly express G2/M- or S-phase markers were annotated as G2/M- or S-phase, respectively. Other cells were annotated as G1 phase cells. Clustering was performed with the “RunUMAP” function in Seurat using significant principal components determined by the JackStraw plot. For each cluster, Marker genes were determined with Seurat’s “FindAllMarkers” function using genes detected in at least 25% of cells and a fold change threshold of 1.8. Sub-clustering of the mesenchymal cluster was performed as above. The developmental processes were calculated as Dorrity *et al*. descripted (Dorrity, 2020).

### Single-cell trajectory reconstruction

The single-cell trajectories were reconstructed using Monocle_ 2.10.1 (Chen et al., 2019, Trapnell et al., 2014, Qiu et al., 2017). A nonlinear reconstruction algorithm, Discriminative Dimensionality Reduction with Trees (DDRTree), was used to reconstruct the single-cell trajectories with genes differentially expressed across four different time points. The state contains cells from E9.5 embryos were set as time zero, and other cells were ordered across the trajectory. Differently expressed genes across pseudotime were selected with a q value less than 0.01. Differential expression analysis between states at branch 1 was performed using the “Beam” function in Monocle. Differently expressed genes were clustered by pseudotime expression patterns to draw the heatmaps.

### Data availability

The single cell RNA-seq datasets have been deposited in the Single-Cell Portal of the Broad Institute under accession number SCP1367.

## Acknowledgments

We are grateful to Drs. Bruce Draper and Lesilee Rose (UC Davis BMCDB program) for their supports, Bridget Mclaughlin and Jonathan Van Dyke (UC Davis Flow Cytometry Shared Resource) for conducting fluorescence-activated cell sorting.

## Additional information

### Funding

| Funder | Grant reference number | Author |
| --- | --- | --- |
| NIH/NIDCR | R01DE026737 | Chengji Zhou |
| NIH/NINDS | R01NS102261 | Chengji Zhou |
| Shriners Hospitals for Children | 85105 | Chengji Zhou |
| The funders had no role in study design, data collection and interpretation, or the decision to submit the work for publication. |  |  |

### Author contributions

YJ, Conception and design, Acquisition of data, Analysis and interpretation of data, Drafting or revising the article; SZ, KR, Acquisition of data, Analysis and interpretation of data, Drafting or revising the article; RG, MM, MI, YL, TI, RD, HZ, DB, Acquisition of data; YX, Analysis and interpretation of data, Drafting or revising the article; CZ, Conception and design, Acquisition of data, Analysis and interpretation of data, Drafting or revising the article.

---

**Figure S1.**
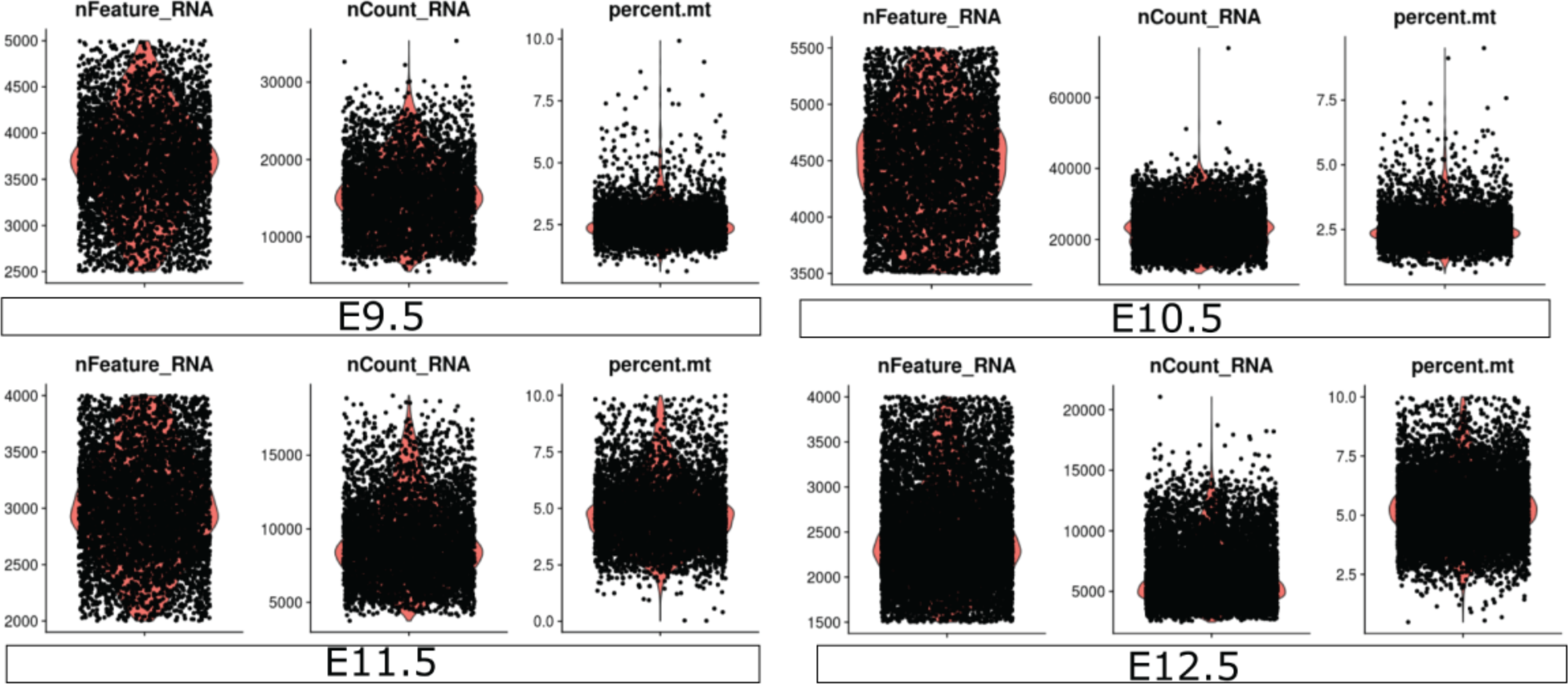
Violin plots for quality control metrics of single cell RNA-seq, including the number of unique molecular identifiers (UMIs) (Left), number of genes (Middle), and percentage of mitochondrial genes (Right) at each age of E9.5∼E12.5.

**Figure S2.**
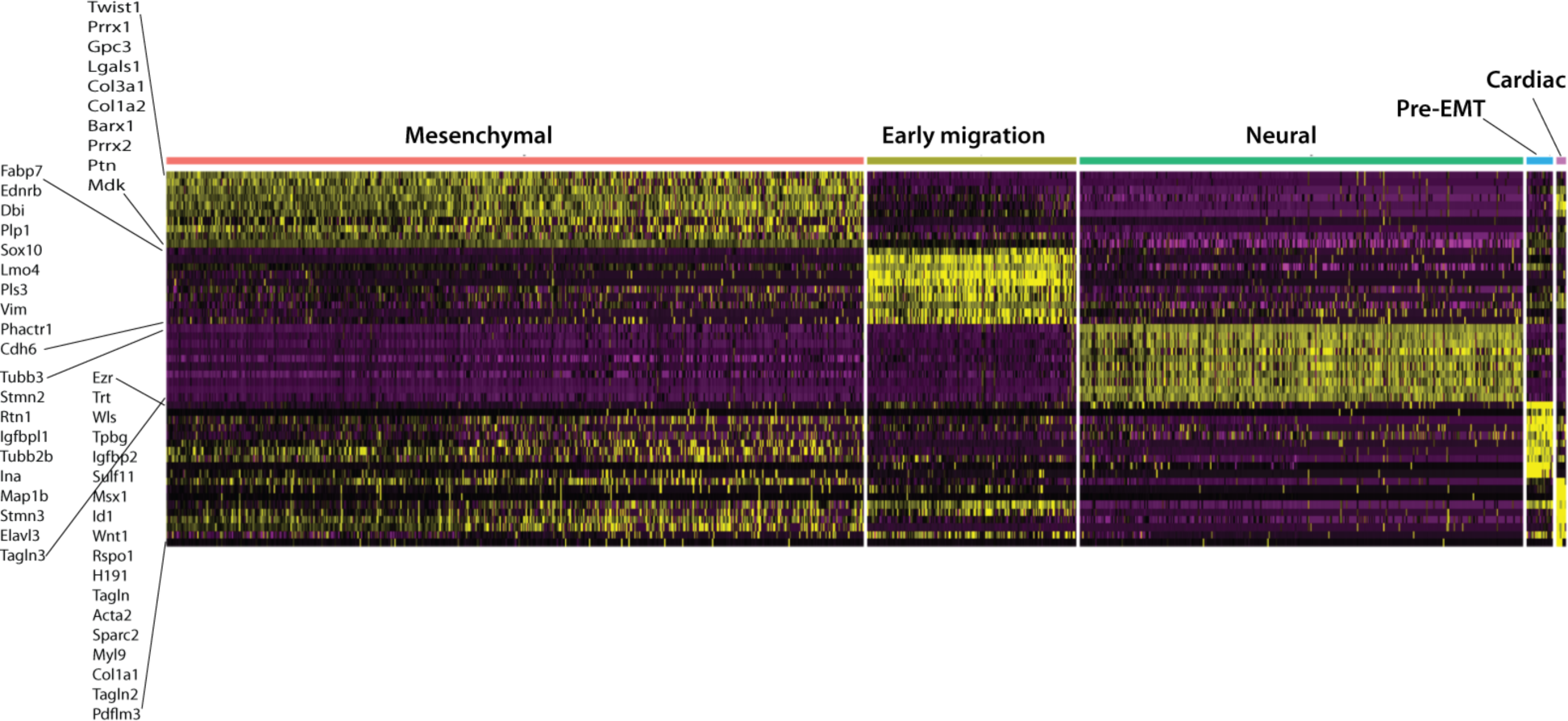
Heatmap showing the expression of the top 10 marker genes for each cell type. The identity of each cluster is labeled on the top of each column.

**Figure S3.**
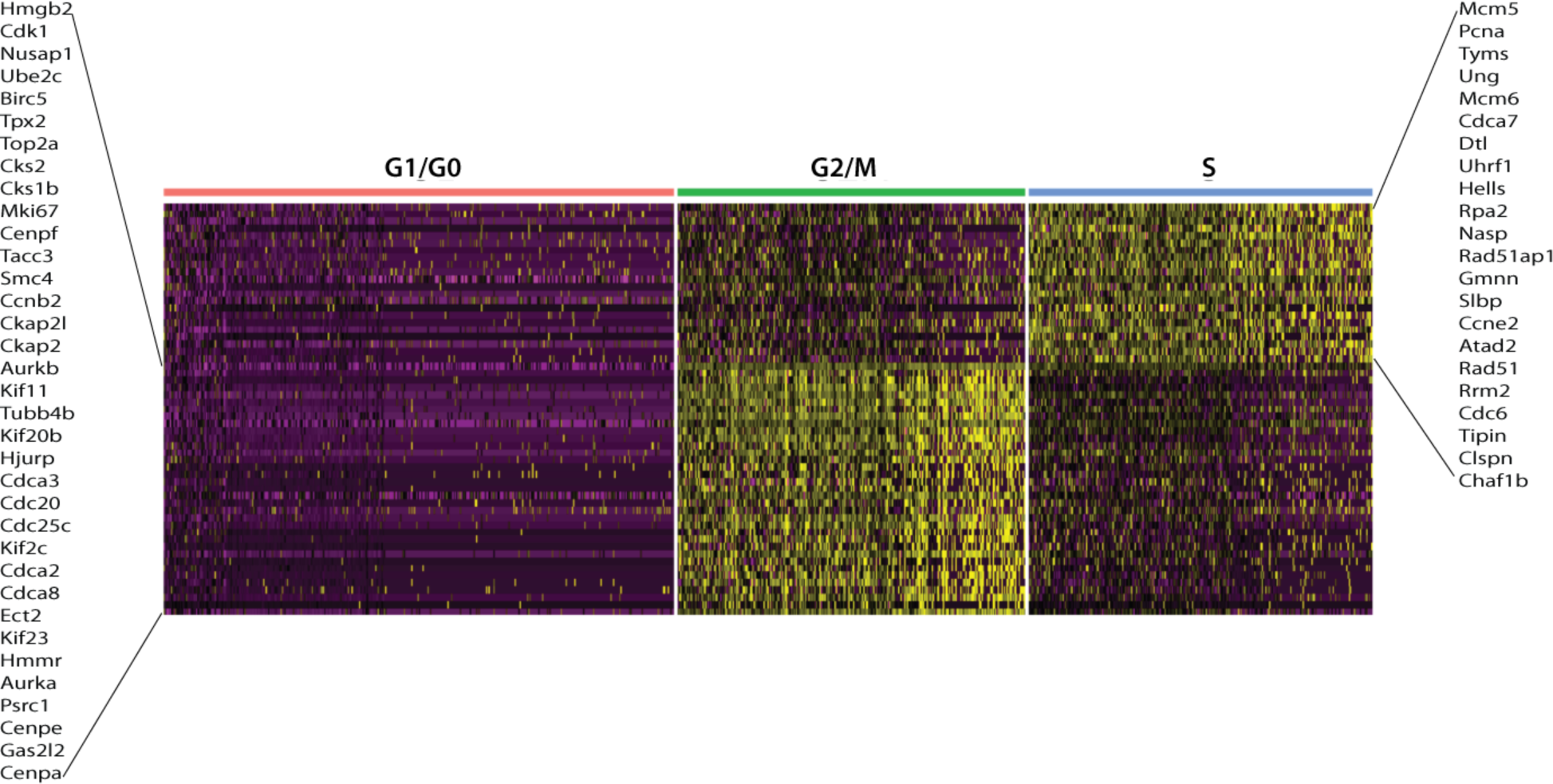
Heatmap of genes used to define the cell cycle phase for each cell.

**Table S1.**
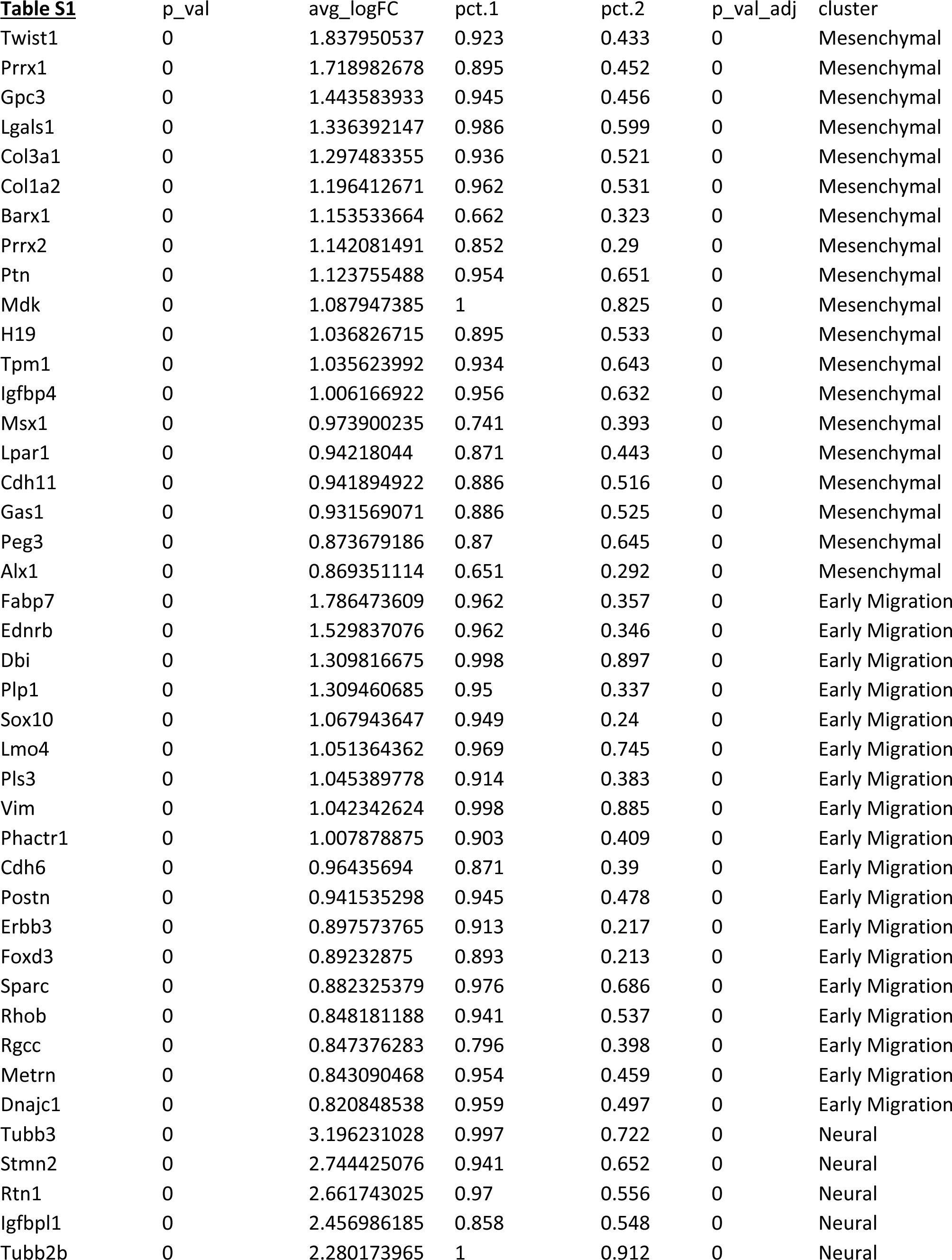
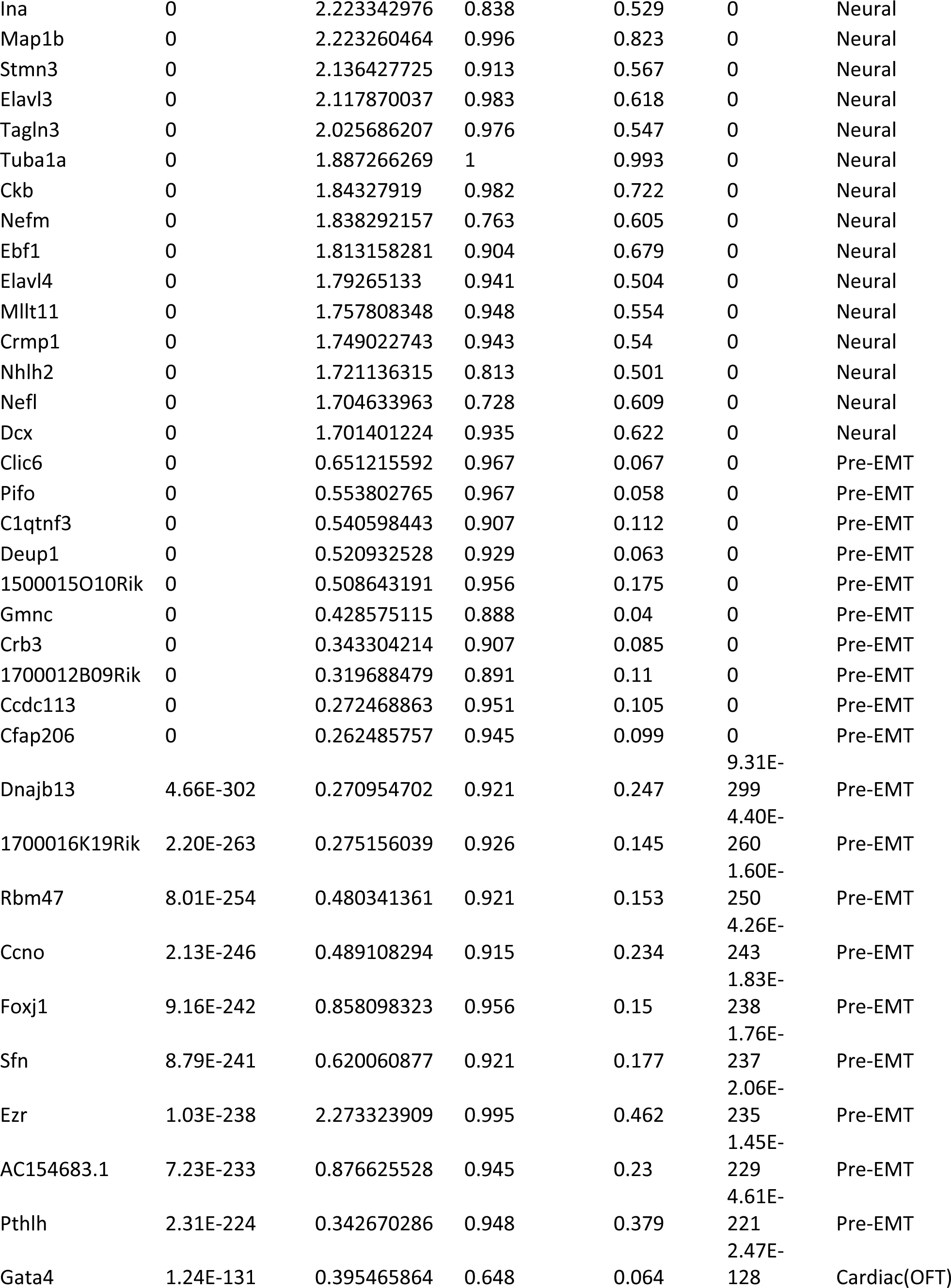
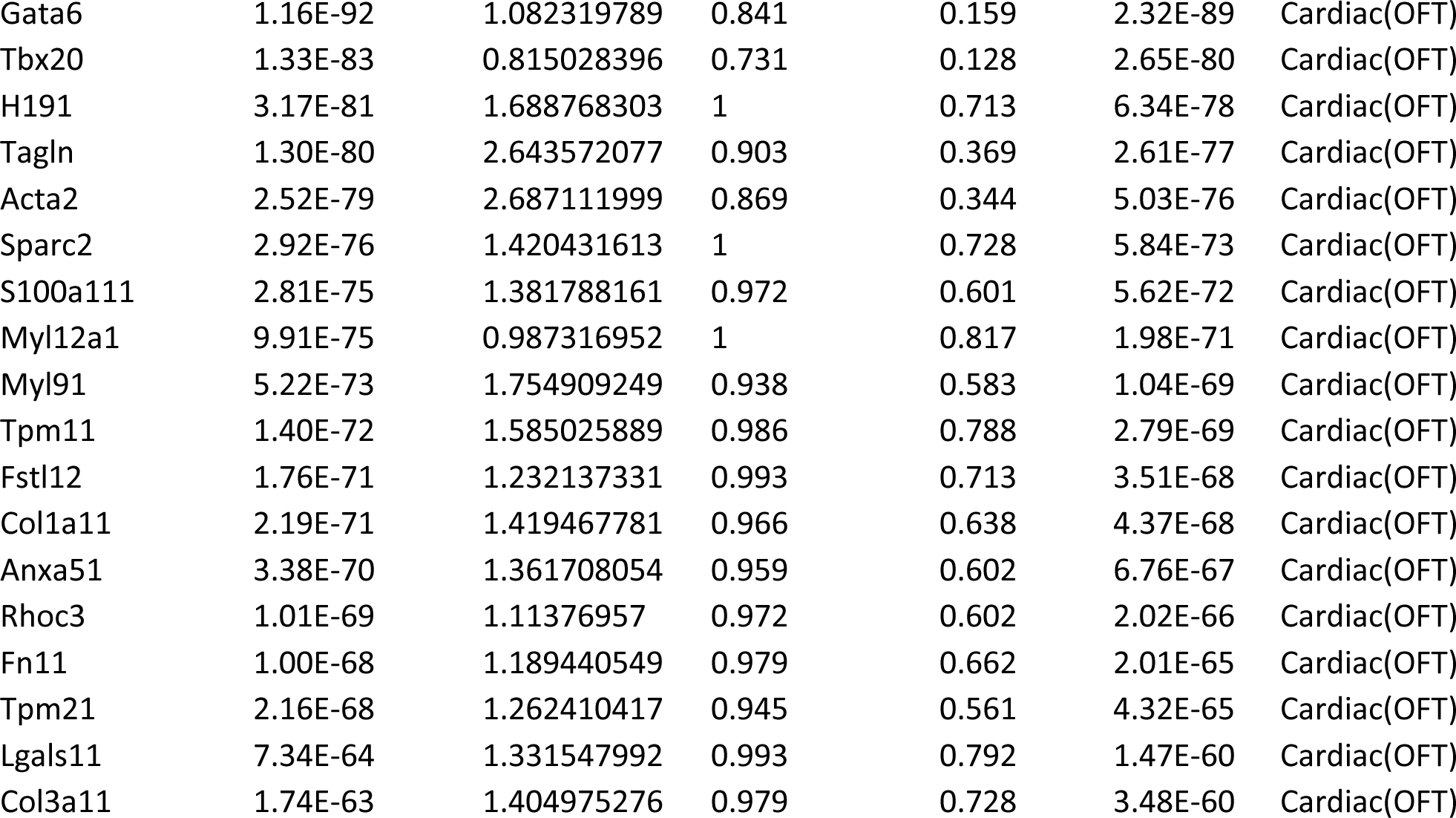
Top marker genes for major NC-derived cell types.

**Table S2.**
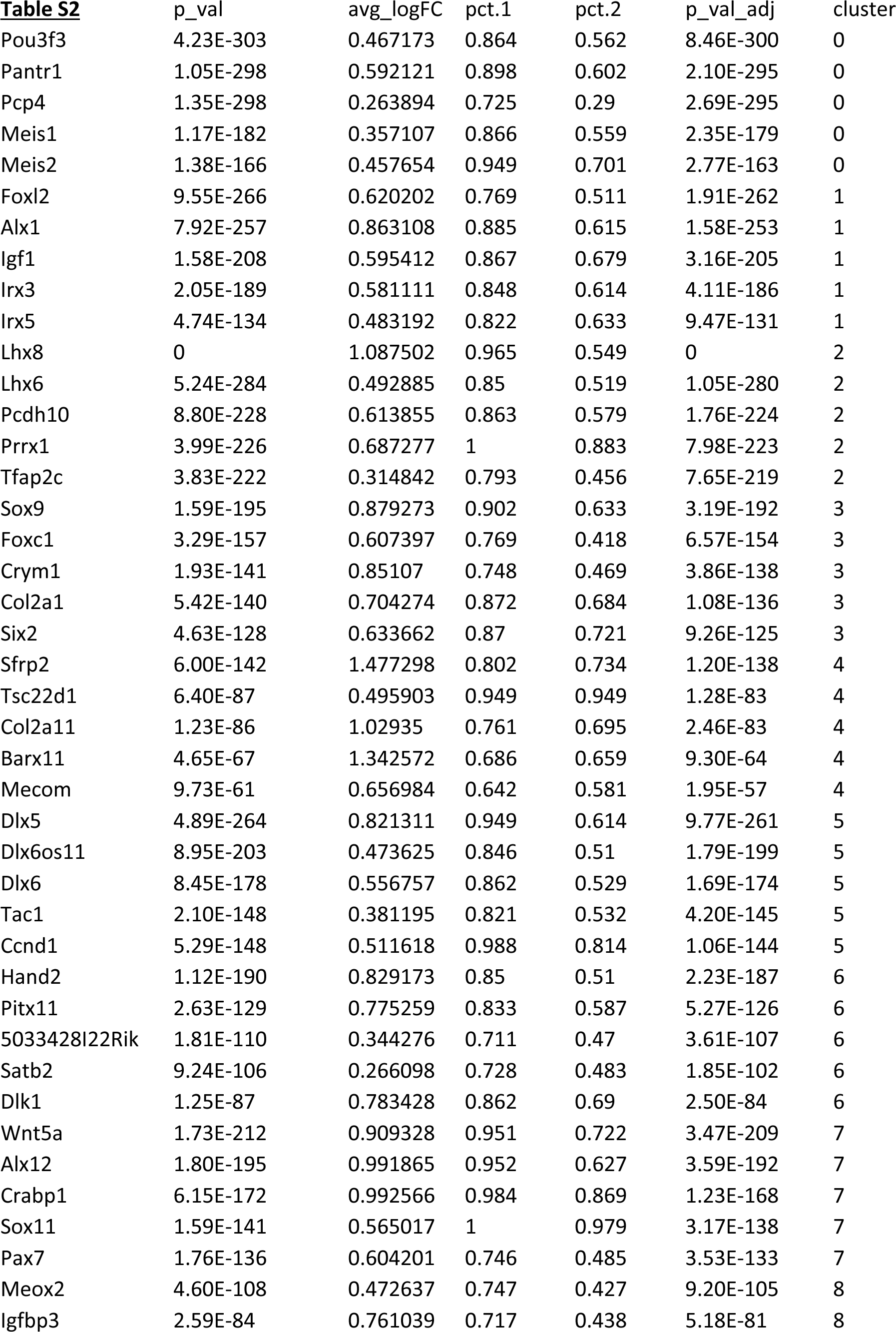
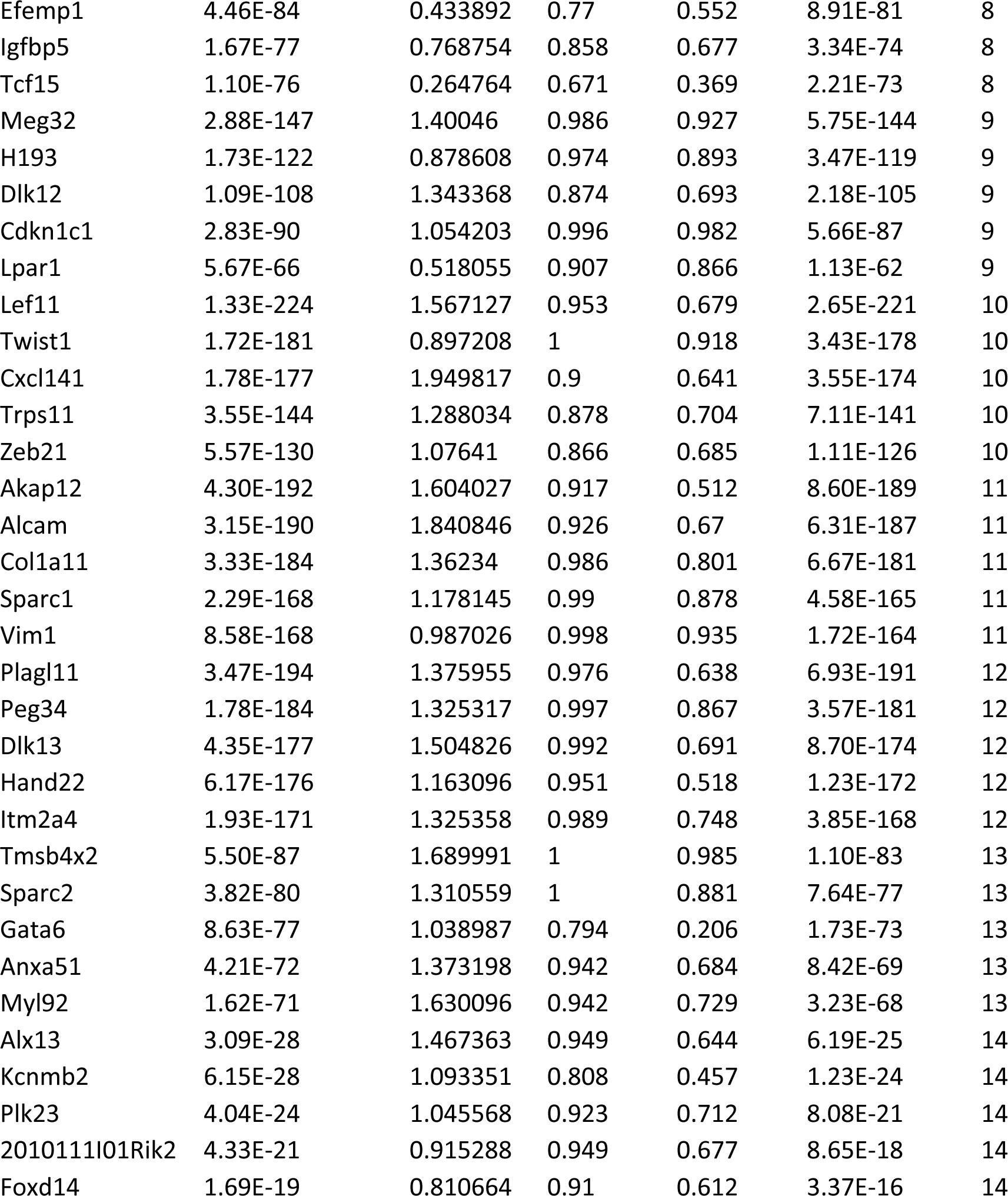
Top marker genes for NC-derived mesenchymal clusters. “p_val” and “p_val_adj” represent the probability for a gene to mark the cluster. avg_logFC, log2 of the ratio of the average expression level of a gene for all cells within the cluster compare to all the other cells. “pct.1” and “pct.2” are the percent of cells in or out the cluster that express the gene.

**Table S3.**
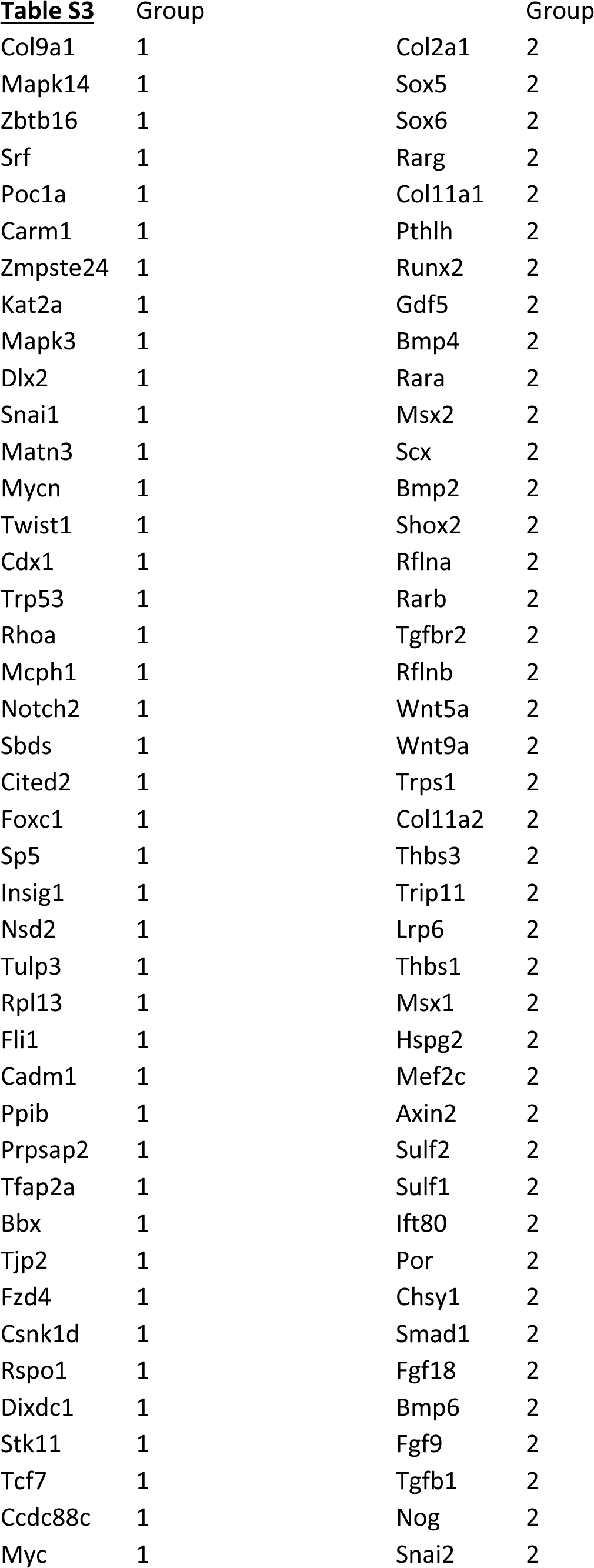
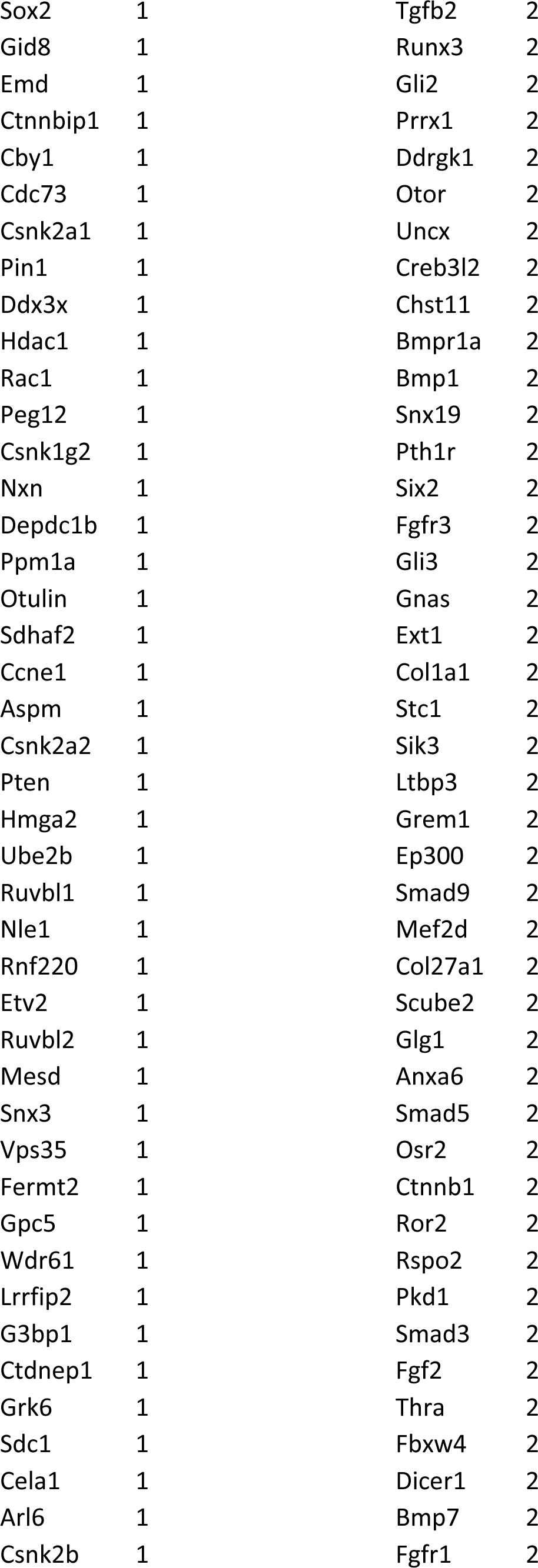
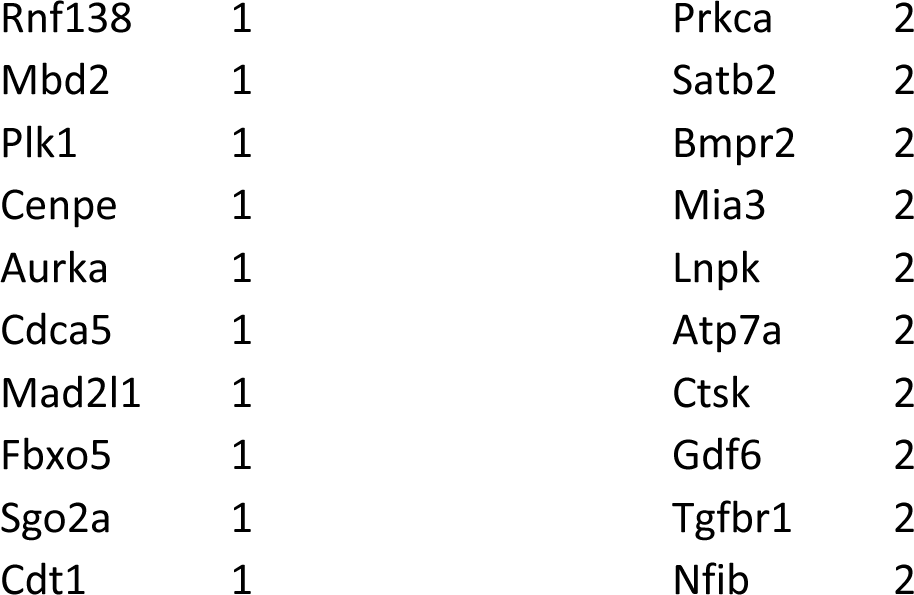
Top gene lists in groups 1 and 2 that exhibit temporal expression patterns, related to figure 5.

**Table S4.** Top gene lists in groups 1∼4 from BEAM Analysis of Branch Point 1, related to **figure 6**.

| **<u>Table S4</u>** | Group |
| --- | --- |
| Crabp1 | 1 |
| Crabp2 | 1 |
| AC160336.1 | 1 |
| Prtg | 1 |
| Ldha | 1 |
| Grrp1 | 1 |
| Srm | 1 |
| Lin28a | 1 |
| Dctpp1 | 1 |
| Aldoa | 1 |
| Rgcc | 2 |
| Fabp7 | 2 |
| Ttyh1 | 2 |
| Car11 | 2 |
| Nell2 | 2 |
| Postn | 2 |
| Gpm6b | 2 |
| Sparc | 2 |
| Zbtb20 | 2 |
| Hey2 | 2 |
| Map1b | 3 |
| Hist3h2ba | 3 |
| Apoe | 3 |
| Nfia | 3 |
| Anxa2 | 3 |
| Ckb | 3 |
| Rspo1 | 3 |
| Scrn1 | 3 |
| Crip1 | 3 |
| Arl4a | 3 |
| Twist1 | 4 |
| Gsta4 | 4 |
| Meis2 | 4 |
| Cd24a | 4 |
| Prrx2 | 4 |
| Slc16a3 | 4 |
| Sfrp2 | 4 |
| Mest | 4 |
| Cited2 | 4 |
| Aprt | 4 |
| Aard | 4 |
| Id1 | 4 |
| Cth | 4 |
| Ptgis | 4 |
| Hacd1 | 4 |
| Naf1 | 4 |

**Table S5.** Top marker genes for E11.5 facial mesenchymal clusters m0∼m3.

|  |  |  |  |  |  |  |
| --- | --- | --- | --- | --- | --- | --- |
| Table S5 | p_val | avg_logFC | pct.1 | pct.2 | p_val_adj | cluster |
| Col2a1 | 3.77E-99 | 1.181788 | 0.856 | 0.467 | 7.53E-96 | m0 |
| Sfrp1 | 7.82E-89 | 0.915514 | 0.861 | 0.519 | 1.56E-85 | m0 |
| Foxc1 | 1.13E-79 | 0.86289 | 0.757 | 0.355 | 2.26E-76 | m0 |
| S100a11 | 5.87E-59 | 0.635084 | 0.96 | 0.867 | 1.17E-55 | m0 |
| Sox9 | 2.12E-58 | 0.997222 | 0.761 | 0.503 | 4.23E-55 | m0 |
| Ebf1 | 6.53E-55 | 0.884611 | 0.82 | 0.584 | 1.31E-51 | m0 |
| Pkdcc | 1.20E-54 | 0.779568 | 0.825 | 0.58 | 2.41E-51 | m0 |
| Six2 | 1.97E-51 | 0.637623 | 0.789 | 0.514 | 3.95E-48 | m0 |
| Zic2 | 5.40E-51 | 0.319418 | 0.427 | 0.066 | 1.08E-47 | m0 |
| Calca | 9.57E-47 | 0.732756 | 0.579 | 0.307 | 1.91E-43 | m0 |
| Alx1 | 1.34E-46 | 0.783765 | 0.908 | 0.64 | 2.68E-43 | m1 |
| Mdk | 2.32E-39 | 0.323551 | 1 | 1 | 4.64E-36 | m1 |
| Mfap4 | 9.50E-25 | 0.386263 | 0.966 | 0.935 | 1.90E-21 | m1 |
| Id3 | 1.31E-22 | 0.421061 | 0.974 | 0.891 | 2.63E-19 | m1 |
| Gata3 | 3.28E-22 | 0.444604 | 0.479 | 0.255 | 6.56E-19 | m1 |
| Tbx2 | 6.13E-22 | 0.407679 | 0.65 | 0.429 | 1.23E-18 | m1 |
| Creb5 | 7.38E-21 | 0.476501 | 0.607 | 0.424 | 1.48E-17 | m1 |
| Mpped2 | 2.76E-19 | 0.462925 | 0.774 | 0.6 | 5.53E-16 | m1 |
| Wnt5a | 2.52E-17 | 0.423862 | 0.762 | 0.62 | 5.03E-14 | m1 |
| Gata2 | 5.39E-17 | 0.366436 | 0.521 | 0.328 | 1.08E-13 | m1 |
| Stmn1 | 5.06E-68 | 0.538458 | 1 | 0.991 | 1.01E-64 | m2 |
| Fabp5 | 2.44E-61 | 0.62328 | 1 | 0.9 | 4.89E-58 | m2 |
| Lix1 | 3.86E-59 | 0.757052 | 0.804 | 0.369 | 7.72E-56 | m2 |
| Crabp1 | 3.07E-56 | 0.871867 | 0.943 | 0.687 | 6.14E-53 | m2 |
| Twist1 | 1.17E-54 | 0.780003 | 0.973 | 0.836 | 2.33E-51 | m2 |
| Lef1 | 2.80E-51 | 0.771265 | 0.714 | 0.394 | 5.61E-48 | m2 |
| Pax1 | 1.19E-50 | 0.6487 | 0.571 | 0.213 | 2.38E-47 | m2 |
| Stmn2 | 7.13E-49 | 0.78273 | 0.557 | 0.204 | 1.43E-45 | m2 |
| Rrm2 | 3.71E-44 | 0.756943 | 0.744 | 0.433 | 7.41E-41 | m2 |
| Al646519 | 7.29E-40 | 0.322885 | 0.473 | 0.13 | 1.46E-36 | m2 |
| Dlk1 | 1.80E-126 | 2.165033 | 0.954 | 0.587 | 3.59E-123 | m3 |
| Cdkn1c | 8.01E-112 | 1.569123 | 1 | 0.989 | 1.60E-108 | m3 |
| Plagl1 | 6.13E-110 | 1.265317 | 0.937 | 0.531 | 1.23E-106 | m3 |
| Mest | 4.34E-108 | 1.079631 | 0.997 | 0.761 | 8.67E-105 | m3 |
| Hand2 | 1.18E-101 | 1.087069 | 0.776 | 0.232 | 2.36E-98 | m3 |
| H19 | 2.63E-95 | 1.118661 | 0.987 | 0.847 | 5.26E-92 | m3 |
| Meg3 | 8.66E-89 | 1.192614 | 0.997 | 0.914 | 1.73E-85 | m3 |
| Peg3 | 4.64E-88 | 1.270328 | 0.967 | 0.789 | 9.27E-85 | m3 |
| Gpc3 | 4.45E-85 | 0.829088 | 0.997 | 0.83 | 8.90E-82 | m3 |
| Nrk | 2.63E-70 | 0.903391 | 0.828 | 0.475 | 5.26E-67 | m3 |

**Table S6.**
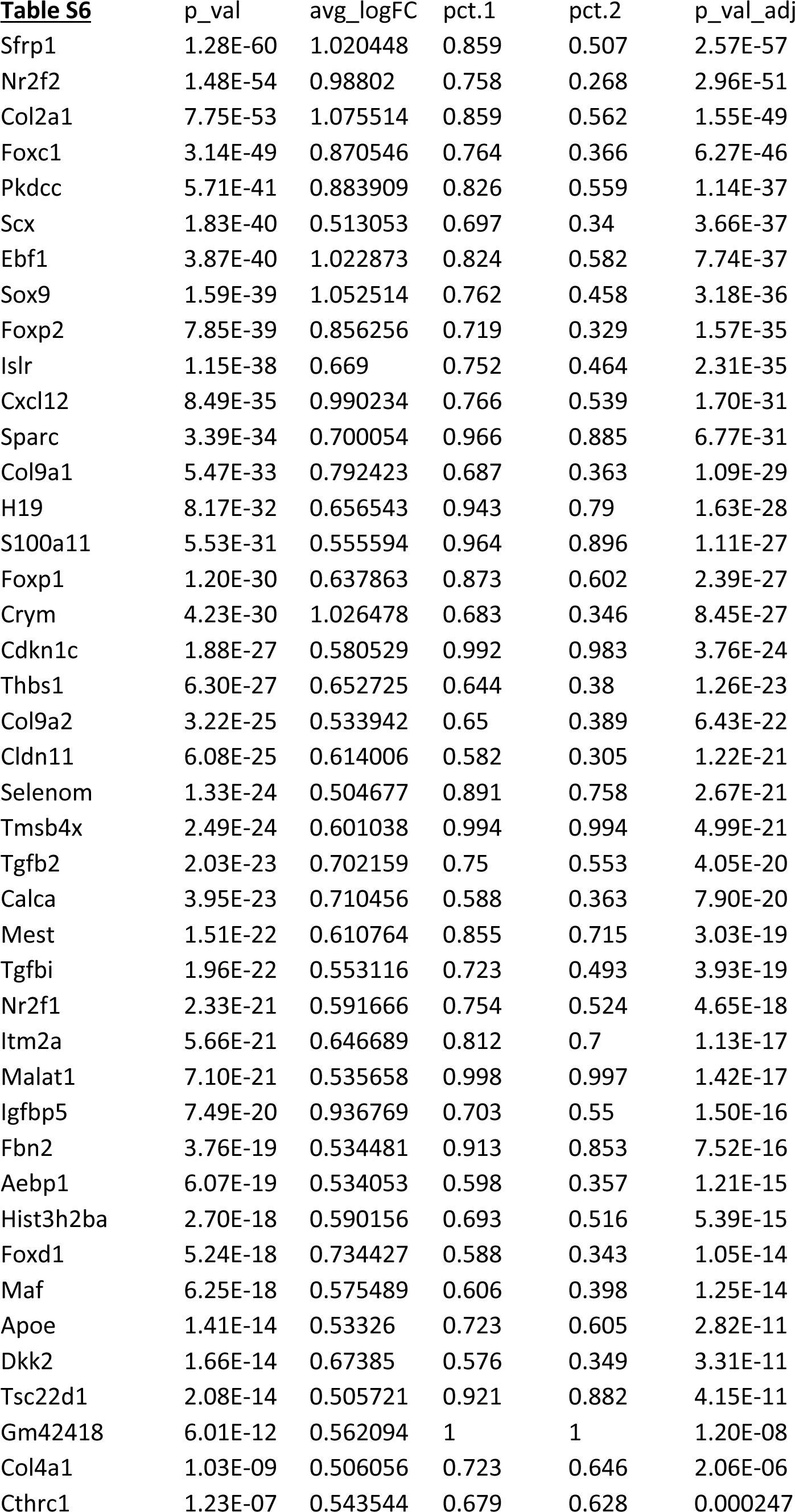
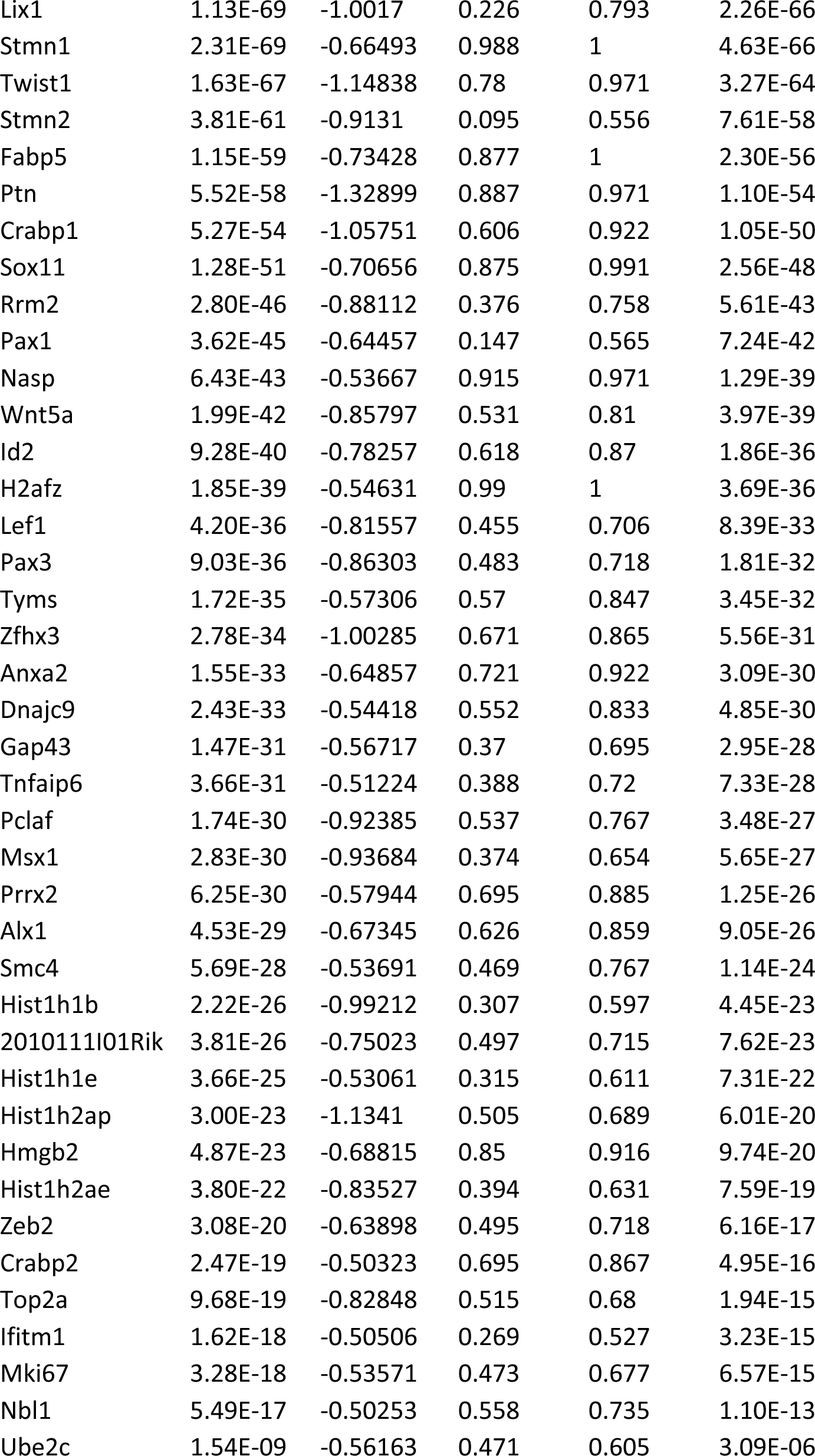
Differentially expressed genes between m0, m2.

**Table S7.**
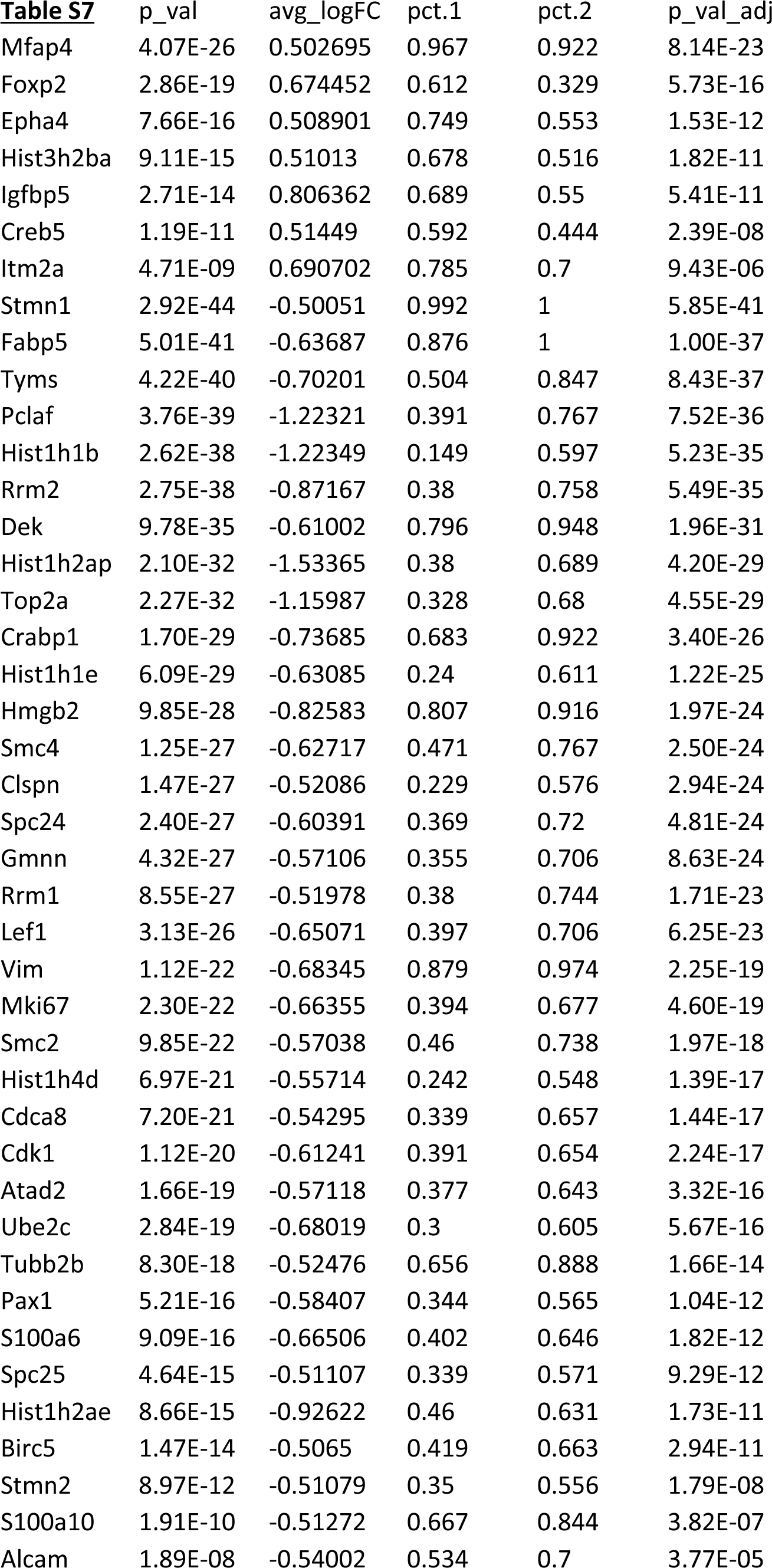
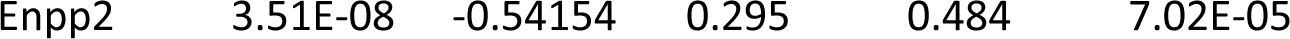
Differentially expressed genes between m1, m2. Highly expressed genes (avg_logFC > 0.2, P adj < 0.05) in m0, m1, or m2 are shown. The percentage of cells that have detected level of gene expression in each cluster (pct.1 and pct.2) are shown.

## Notes

### Competing Interest Statement

The authors have declared no competing interest.

